# Modeling the cell-type specific mesoscale murine connectome with anterograde tracing experiments

**DOI:** 10.1101/2023.05.02.539079

**Authors:** Samson Koelle, Dana Mastrovito, Jennifer D Whitesell, Karla E Hirokawa, Hongkui Zeng, Marina Meila, Julie A Harris, Stefan Mihalas

**Affiliations:** Allen Institute for Brain Science, Seattle, WA, USA; Department of Statistics, University of Washington, Seattle, WA, USA

**Keywords:** Connectivity, Cell-type, Mouse

## Abstract

The Allen Mouse Brain Connectivity Atlas (MCA) consists of anterograde tracing experiments targeting diverse structures and classes of projecting neurons. Beyond regional anterograde tracing done in C57BL/6 wild type mice, a large fraction of experiments are performed using transgenic Cre-lines. This allows access to cell-class specific whole brain connectivity information, with class defined by the transgenic lines. However, even though the number of experiments is large, it does not come close to covering all existing cell classes in every area where they exist. Here, we study how much we can fill in these gaps and estimate the cell-class specific connectivity function given the simplifying assumptions that nearby voxels have smoothly varying projections, but that these projection tensors can change sharply depending on the region and class of the projecting cells.

This paper describes the conversion of Cre-line tracer experiments into class-specific connectivity matrices representing the connection strengths between source and target structures. We introduce and validate a novel statistical model for creation of connectivity matrices. We extend the Nadaraya-Watson kernel learning method which we previously used to fill in spatial gaps to also fill in a gaps in cell-class connectivity information. To do this, we construct a "cell-class space" based on class-specific averaged regionalized projections and combine smoothing in 3D space as well as in this abstract space to share information between similar neuron classes. Using this method we construct a set of connectivity matrices using multiple levels of resolution at which discontinuities in connectivity are assumed. We show that the connectivities obtained from this model display expected cell-type and structure specific connectivities. We also show that the wild type connectivity matrix can be factored using a sparse set of factors, and analyze the informativeness of this latent variable model.

**AUTHOR SUMMARY:** Large-scale studies have described the connections between areas in multiple mammalian models in ever expanding detail. Standard connectivity studies focus on the connection strength between areas. However, when describing functions at a local circuit level, there is an increasing focus on cell types. We have recently described the importance of connection types in the cortico-thalamic system, which allows an unsupervised discovery of its hierarchical organization. In this study we focus on adding a dimension of connection type for a brain-wide mesoscopic connectivity model. Even with our relatively massive dataset, the data in the cell type direction for connectivity is quite sparse, and we had to develop methods to more reliably extrapolate in such directions, and to estimate when such extrapolations are impossible. This allows us to fill in such a connection type specific inter-areal connectivity matrix to the extent our data allows. While analyzing this complex connectivity, we observed that it can be described via a small set of factors. While not complete, this connectivity matrix represents a a categorical and quantitative improvement in mouse mesoscale connectivity models.

## 1 INTRODUCTION

The mammalian nervous system enables an extraordinary range of natural behaviors, and has inspired much of modern artificial intelligence. Neural connections including those from one region to another form the architecture underlying this capability. These connectivities vary by neuron type, as well as source (cell body) location and target (axonal projection) structures. Thus, characterization of the relationship between neuron type and source and target structure is important for understanding the overall nervous system.

Viral tracing experiments - in which a viral vector expressing GFP is transduced into neural cells through stereotaxic injection - are a useful tool for mapping these connections on the mesoscale (Chamberlin, Du, de Lacalle, & Saper, 1998; Daigle et al., 2018; J. A. Harris, Oh, & Zeng, 2012). The GFP protein moves into the axon of the projecting neurons. The long range connections between different areas are generally formed by axons which travel from one region to another. Two-photon tomography imaging can be used to determine the location and strength of the fluorescent signals in two-dimensional slices. These locations can then be mapped back into three-dimensional space, and the signal may then be integrated over area into cubic voxels to give a finely-quantized three-dimensional fluorescence.

Several statistical models for the conversion of such experiment-specific signals into generalized estimates of connectivity strength have been proposed (Gămănut et al., 2018; K. D. Harris, Mihalas, & Shea-Brown, 2016; Knox et al., 2019; Oh et al., 2014). Of these, Oh et al. (2014) and Knox et al. (2019) provide a model for **regionalized connectivities**, which are voxel connectivities integrated by region. The value of these models is that they provide some improvement over simply averaging the projection signals of injections in a given region. However, these previous works only model connectivities observed in wild type mice which are suboptimally suited to assessment of cell-type specific connectivity compared with fluorescence from Cre-recombinase induced eGFP expression in cell-types specified by the combination of transgenic mouse strain and transgene promoter (J. A. Harris et al., 2019). We generally refer to sets of targeted eGFP-expressing cells in tracing experiments as a **cell class** since they may contain multiple types. For example, use of both wild-type and transgenic mice would give rise to cell-class specific experiments, albeit with different yet perhaps overlapping classes of cells.

Thus, this paper introduces a class-specific statistical model for anterograde tracing experiments that synthesizes the diverse set of **Cre-lines** described in J. A. Harris et al. (2019), and expands this model to the entire mouse brain. Our model is a to-our-knowledge novel statistical estimator that takes into account both the spatial position of the labelled source, as well as the categorical cell class. Like the previously state-of-the-art model in Knox et al. (2019), this model predicts regionalized connectivity as an average over positions within the structure, with nearby experiments given more weight. However, our model weighs class-specific behavior in a particular structure against spatial position, so a nearby experiment specific to a similar cell-class is relatively up-weighted, while a nearby experiment specific to a dissimilar class is down-weighted. This model outperforms the model of Knox et al. (2019) based on its ability to predict held-out experiments in leave-one-out cross-validation. We use the trained model to estimate overall connectivity matrices for each assayed cell class.

The resulting cell-type specific connectivity is a directed weighted multigraph which can be represented as a tensor with missing values. We do not give an exhaustive analysis of this data, but do establish a lower-limit of detection, verify several cell-type specific connectivity patterns found elsewhere in the literature, and show that these cell-type specific signals are behaving in expected ways. We also decompose the wild type connectivity matrix into factors representing latent connectivity patterns, which we call archetypes. These components allow approximation of the regionalized connectivity using linear combinations of a small set of components.

Section 2 gives information on the data and statistical methodology, and Section 3 presents our results. These include connectivities, assessments of model fit, and subsequent biological and statistical analyses. Additional information on our dataset, methods, and results are given in Supplemental Sections 5, 6, and 7, respectively.

## 2 METHODS

We estimate and analyze cell class-specific connectivity functions using models trained on murine brain viral tracing experiments. This section describes the data used to generate the model, the model itself, the evaluation of the model against its alternatives, and the use of the model in creation of the connectivity estimate matrices. It also includes background on the non-negative matrix factorization method used for decomposing the wild type connectivity matrix into latent factors. Additional information about our data and methods are given in Supplemental Sections 5 and 6, respectively.

### Data

Our dataset *𝒟* consists of *n =* 1751 publicly available murine brain viral tracing experiments from the Allen Mouse Brain Connectivity Atlas. Figure 1a summarizes the experimental process used to generate this data. In each experiment, a mouse is injected with an adeno-associated virus (AAV) encoding green fluorescent protein (GFP) into a single location in the brain. Location of fluorescence is mediated by the location of the injection, the characteristics of the transgene, and the genotype of the mouse. In particular, Cre-driver or, equivalently, Cre-line mice are engineered to express Cre under the control of a specific and single gene promoter. This localizes expression of Cre to regions with certain transcriptomic cell-types signatures. In such Cre-driver mice, we used a double-inverted floxed AAV to produce eGFP fluorescence that depends on Cre expression in infected cells. To account for the complex cell-type targeting induced by a particular combination of Cre-driver genotype and GFP promoter, we refer to the combinations of cell-types targeted by a particular combination of AAV and Cre-driver mice as cell-classes. For example, we include experiments from Cre-driver lines that selectively label cell classes located in distinct cortical layers or other nuclei across the whole brain. For injections in the wild type mice, we used the Synapsin I promoter (Jackson, Dayton, Deverman, & Klein, 2016; Kügler, Kilic, & Bähr, 2003). For injections into Cre mice, we used the CAG promoter with a Flex cassette for Cre-mediated recombination control (Saunders, Johnson, & Sabatini, 2012). Additional details on are given in J. A. Harris et al. (2019).

**Figure 1:**
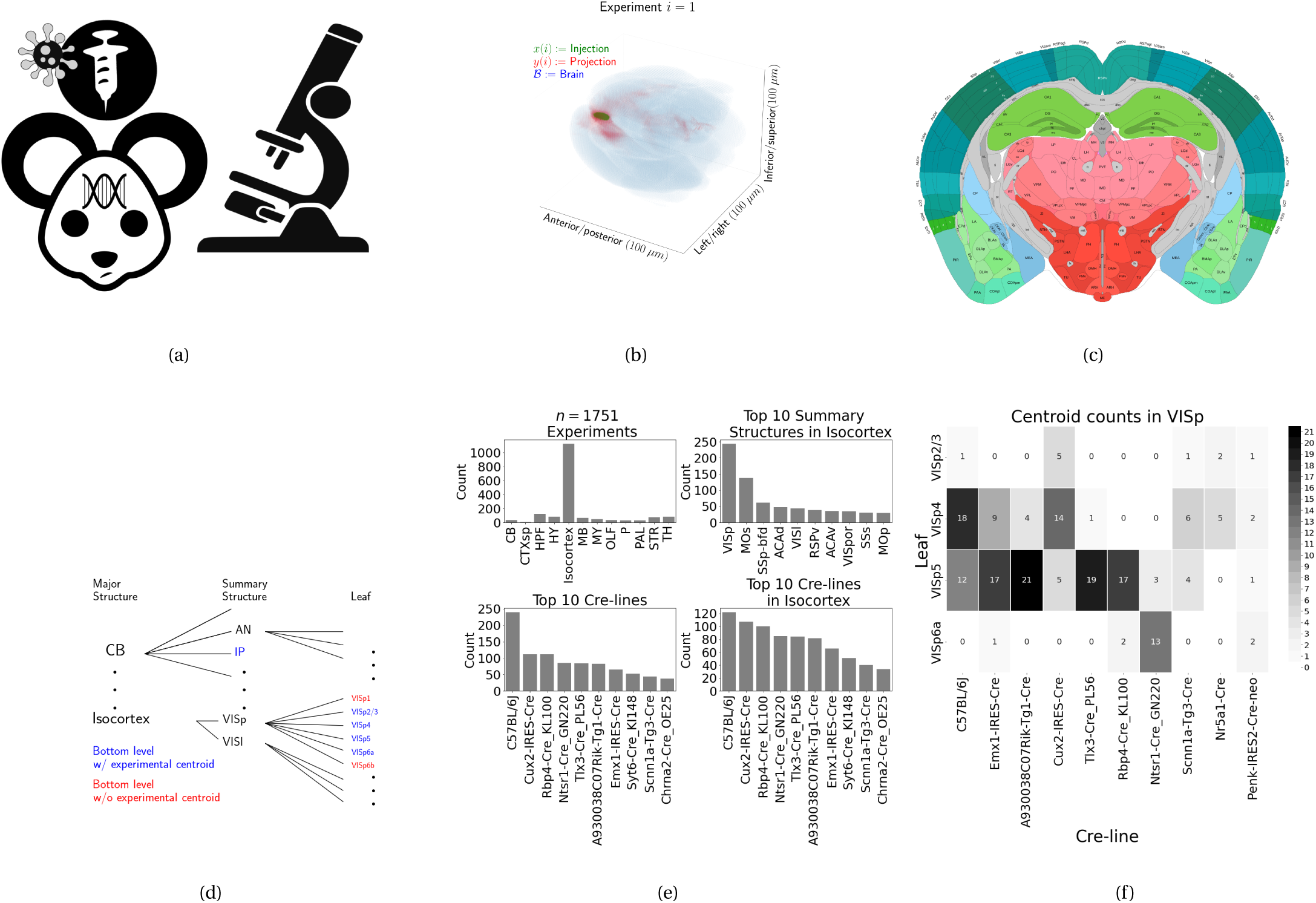
Experimental setting. 1a For each experiment, a Cre-dependent eGFP-expressing transgene casette is transduced by stereotaxic injection into a Cre-driver mouse, followed by serial two-photon tomography imaging. 1b An example of the segmentation of projection (targets) and injection (source) for a single experiment. Within each brain (blue), injection (green) and projection (red) areas are determined via histological analysis and alignment to the Allen Common Coordinate Framework (CCF). 1c Brain region parcellations within a coronal plane of CCFv3. 1d Explanation of nested structural ontology highlighting various levels of CCFv3 structure ontology. Lowest-level (leaf) structures are colored in blue, and structures without an injection centroid are colored in red. 1e Abundances of tracer experiments by Cre-line and region of injection. 1f Co-occurrence of layer-specific centroids and Cre-lines within VISp.

For each experiment, the fluorescent signal imaged after injection is aligned into the Allen Common Coordinate Framework (CCF) v3, a three-dimensional average template brain that is fully annotated with regional parcellations Wang et al. (2020). The whole brain imaging and registration procedures described in detail in Kuan et al. (2015); Oh et al. (2014) produce quantitative metrics of fluorescence discretized at the 100 *µ*m **voxel** level. Given an experiment, this image was histologically segmented by an analyst into *injection* and *projection* areas corresponding to areas containing somas, dendrites and axons or exclusively axons of the transfected neurons. An example of a single experiment rendered in 3D is given in Figure 1b. Given an experiment *i*, we represent injections and projections as functions *x*(*i*), *y* (*i*): *ℬ →* **ℝ***_≥_*_0_, where *ℬ* ⊂ [1 : 132] *×* [1 : 80] *×* [1 : 104] corresponds to the subset of the (1.32 *×* 0.8 *×* 1.04) cm rectangular space occupied by the standard voxelized mouse brain. We also calculate injection centroids *c*(*i*) ∈ **ℝ**^3^ and regionalized projections *y_𝒯_* (*i*) ∈ **ℝ***^T^* given by the sum of *y* (*i*) in each region. A description of these steps is in Supplemental Section 6.

Our goal is the estimation of **regionalized connectivity** from one region to another. A visual depiction of this region parcellation for a two-dimensional slice of the brain is given in Figure 1c. All structures annotated in the CCF belong to a hierarchically ordered ontology, with different areas of the brain are parcellated to differing finer depths within a hierarchical tree. We denote the main levels of interest as major structures, summary structures, and layers. Not every summary structure has a layer decomposition within this ontology, so we typically consider the finest possible regionalization - for example, layer within the cortex, and summary structure within the thalamus, and denote these structures as leafs. As indicated in Figure 1d, the dataset used to generate the connectivity model reported in this paper contains certain combinations of region and cell class frequently, and others not at all. A summary of the most frequently assayed cell classes and structures is given in Figures 1e and 1f. Since users of the connectivity matrices may be interested in particular combinations, or interested in the amount of data used to generate a particular connectivity estimate, we present this information about all experiments in Supplemental Section 5.

### Modeling Regionalized Connectivity

We define voxelized cell-class specific connectivity *f* : *𝒱 × ℬ ×* → **ℝ***_≥_*_0_ as giving the voxelized connectivity strength of a particular cell class from a source voxel to a target voxel. In contrast to Knox et al. (2019), which only uses wild type C57BL/6J mice, our dataset has experiments targeting *|𝒱 |=* 114 different combinations of Cre-driver mice and Cre-regulated AAV transgenes jointly denoted as *𝒱* :*=* {*v* }. As in Knox et al. (2019), we ultimately estimate an integrated regionalized connectivity defined with respect to a set of *S =* 564 source leafs *ℓ* :*=* {*s*} and *T =* 1123 target leafs *𝒯* :*=* {*t* }, of which 1123−564 *=* 559 are contralateral. That is, we define

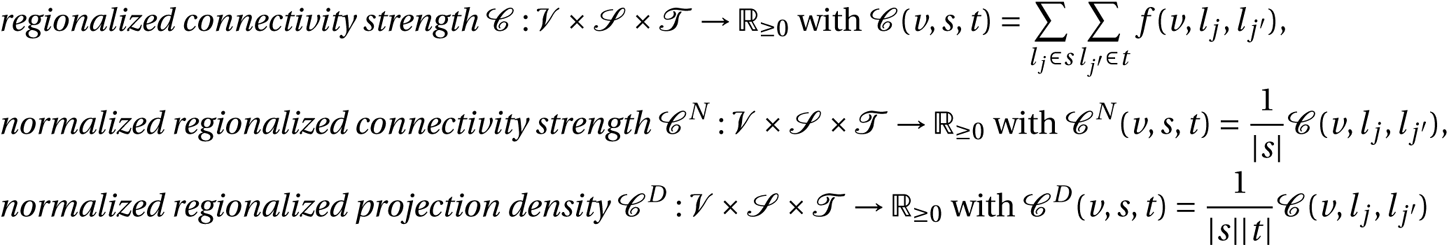

where *l _j_* and *l _j_’* are the locations of source and target voxels, and *|s|* and *|v|* are defined to be the number of voxels in the source and target structure, respectively. Since the normalized strength and densities are computable from the strength via a fixed normalization, our main statistical goal is to estimate 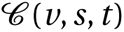 for all *v*, *s* and *t* .In other words, we want to estimate matrices 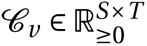. We call this estimator 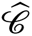.

Construction of such an estimator raises the questions of what data to use for estimating which connectivity, how to featurize the dataset, what statistical estimator to use, and how to reconstruct the connectivity using the chosen estimator. We represent these considerations as

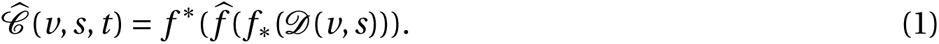

This makes explicit the data featurization *f_*_*, statistical estimator 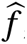, and any potential subsequent transformation *f \** such as summing over the source and target regions. Denoting *𝒟* as a function of *v* and *s* reflects that we consider using different data to estimate connectivities for different cell-classes and source regions. Table 1 reviews estimators used for this data-type used in previous work, as well as our two main extensions: the Cre-NW and **Expected Loss** (EL) models. The main differences in our data featurization from (Knox et al., 2019) are that we regionalize our data at the leaf level where available so that it layer-specific behavior is visible, and normalize our data by projection signal in order to account for differences between cell class. Additional model selection results are given in Supplemental Section 5 for alternative normalization strategies, and more detail on estimation is given in Supplemental Section 6.

**Table 1:** Estimation of *𝒷* using connectivity data. The regionalization, estimation, and featurization steps are denoted by *f \**, 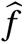, and *f_*_*, respectively. The training data used to fit the model is given by index set *I* . We denote experiments with centroids in particular major brain divisions and leafs as *I_m_* and *I_l_*, respectively. Data *I_l_* /*I_m_* means that, given a location *l_s_* ∈ *s* ∈ *m*, the model 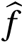 is trained on all of *I_m_*, but only uses *I_l_* for prediction. The non-negative least squares estimator (NNLS) fits a linear model that predicts regionalized projection signal *y_𝒯_* as a function of regionalized injection signal *x_*ℓ*_* . Thus, the regionalization step for a region *s* is given by applying the learned matrix 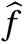to the *s*-th indicator vector. In contrast, the Nadaraya-Watson model (NW) is a local smoothing model that generates a prediction for each voxel within the source structure that are then averaged to create estimate the structure-specific connectivity.

| Name | $f^*$ | $\hat{f}$ | $f_*$ | $\mathcal{D}(\nu, s)$ |
| --- | --- | --- | --- | --- |
| NNLS (Oh et al., 2014) | $\hat{f}(1_s)$ | NNLS(X,Y) | $X = x_{\mathcal{S}}, Y = y_{\mathcal{T}}$ | $I_m / I_m$ |
| NW (Knox et al., 2019) | $\sum_{l_s \in s} \hat{f}(l_s)$ | NW(X,Y) | $X = l_s, Y = y_{\mathcal{T}}$ | $I_m / I_m$ |
| Cre-NW | $\sum_{l_s \in s} \hat{f}(l_s)$ | NW(X,Y) | $X = l_s, Y = y_{\mathcal{T}}$ | $(I_l \cap I_\nu) / I_m$ |
| Expected Loss (EL) | $\sum_{l_s \in s} \hat{f}(s)$ | EL(X, Y, $\nu$) | $X = l_s, Y = y_{\mathcal{T}}, \nu$ | $I_l / I_m$ |

Our contributions - the Cre-NW and Expected Loss (EL) models - have several differences from the previous methods. In contrast to the non-negative least squares (Oh et al., 2014) and Nadaraya-Watson (Knox et al., 2019) estimators that account only for source region *s*, our new estimators account cell class *v*, The Cre-NW estimator only uses experiments from a particular class to predict connectivity for that class, while the EL estimator shares information between classes within a structure. Both of these estimator take into account both the cell-class and the centroid position of the experimental injection. Like the NW and Cre-NW estimator, the EL estimator generates predictions for each voxel in a structure, and then sums them together to get the overall connectivity. However, in contrast to the NW approaches, the EL estimate of the projection vector for a cell-class at a location weights the average projection of that cell-class in the region containing the location against the relative locations of all experimental centroids in the region regardless of class. That is, cell-class and source region combinations with similar average projection vectors will be upweighted when estimating 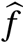. Thus, all experiments that are nearby in three-dimensional space can help generate the prediction, even when there are few nearby experiments for the cell-class in question. A detailed mathematical description of our new estimator is given in Supplemental Section 6.

### Model evaluation

We select optimum functions from within and between our estimator classes using **leave-one-out cross validation**, in which the accuracy of the model is assessed by its ability to predict projection vectors experiments excluded from the training data on the basis of their cell class and experimental centroid. Equation 1 includes a deterministic step *f \** included without input by the data. The performance of 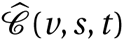 is thus determined by performance of 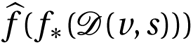. Thus, we evaluate prediction of 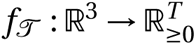 the regionalized connection strength at a given location.

Another question is what combinations of *v*, *s*, and *t* to generate a prediction for. Our EL and Cre-NW models are leaf specific. They only generate predictions for cell-classes in leafs where at least one experiment with a Cre-line targeting that class has a centroid. To accurately compare our new estimators with less-restrictive models such as used in Knox et al. (2019), we restrict our evaluation set to Cre driver/leaf combinations that are present at least twice. The sizes of these evaluation sets are given in Supplemental Section 5.

We use weighted *l* 2-loss to evaluate these predictions.

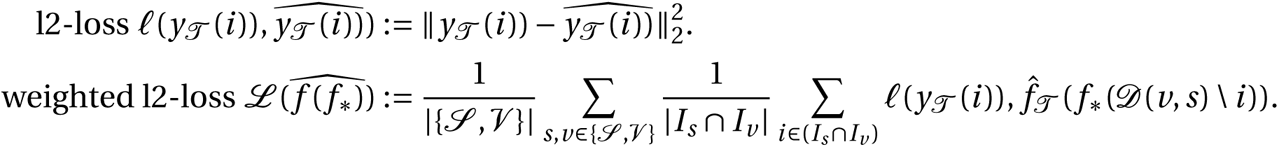

*I_s_* refers to the set of experiments with centroid in structure *s*, and *I_v_* refers to the set of experiments with Cre-line *v*, so *|I_s_* ∩*I_v_ |* is the number of experiments of Cre-line *v* with injection centroid in structure *s*. This is a somewhat different loss from Knox et al. (2019) because of the increased weighting of rarer combinations of *s* and *v* implicit in the 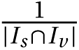 term in the loss. The establishment of a lower limit of detection and the extra cross-validation step used in the EL model to establish the relative importance of regionally averaged cell-class projection and injection centroid position are covered in Supplemental Section 6.

### Connectivity analyses

We examine our connectome estimates with both comparisons to known biology and statistical decompositions. As an exploratory analysis, we use heirarchical clustering to compare outputs from connectivities from different Cre-lines. Details of and results from this approach are given in Section 7. We then use non-negative matrix factorization (NMF) to factor the wild-type connectivity matrix into a small set of underlying components that can be linearly combined to reproduce the observed long-range wild-type connectivity. Inspired by Mohammadi, Ravindra, Gleich, and Grama (2018), we refer to these latent coordinates as **connectivity archetypes** since they represent underlying patterns from which we can reconstruct a broad range of observed connectivities, although we note that the genomic archetypal analysis in that paper is slightly methodologically distinct.

NMF refers to a collection of **dictionary-learning** algorithms for decomposing a non-negatively-valued matrix such as *𝒷* into positively-valued matrices called, by convention, weights 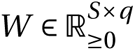 and hidden units 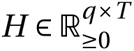. NMF assumes a simple linear statistical model: that the observed matrix is composed of linear combinations of latent coordinates (Devarajan, 2008). Unlike PCA, NMF specifically accounts for the fact that data are all in the positive orthant, and it is more stable and interpretable in assays of complex biological systems than heirarchical clustering (Brunet, Tamayo, Golub, & Mesirov, 2004) The choice of matrix factorization method reflects particular scientific subquestions and probabilistic interpretations, and the matrix *H* may used to identify latent structures with interpretable biological meaning.

Our application of NMF to decompose the estimated long-range connectivity is some independent interest, since we ignore connections between source and target regions less than 1500 *µm* apart. This is because short-range projections resulting from diffusion and traveling fibers dominate the matrices 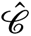. Our NMF algorithm solves the following optimization problem

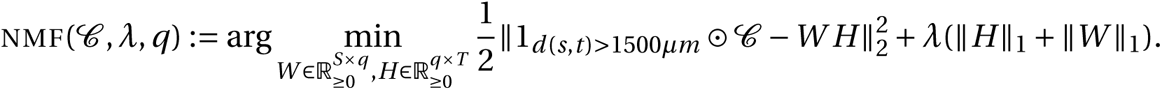

We set *λ* = 0.002 to encourage sparser and therefore more interpretable components. We use unsupervised cross-validation to determine an optimum *q*, and show the top 15 stable components (Perry, 2009). Since the NMF objective is difficult to optimize and sensitive to initialization, we follow up with a stability analysis via clustering the resultant *H* from multiple replicates. The medians of the component clusters appearing frequently across NMF replicates are selected as **connectivity archetypes**. Details of these approaches are given in Supplementary Sections 6 and 7.

## 3 RESULTS

The main result of this paper is the creation of cell-type specific connectivity estimates from the Allen Mouse Brain Connectivity Atlas (MCA) experiments. We first establish that our new expected-loss (EL) estimator performs best in validation assays for estimating wild-type and cell-type specific connectivities. We then show that Cre-specific connectivity matrices generated using this model are consistent with known biology. Finally, we factor some of these connectivity matrices to show how connectivity arises from latent components, and that these latent components may be associated with cell-types.

### Model evaluation

Our EL model generally performs better than the other estimators that we consider. Table 2a contains weighted losses from leave-one-out cross-validation of candidate models, such as the NW Major-WT model from Knox et al. (2019). The EL model combines the good performance of class-specific models like NW Leaf-Cre in regions like Isocortex with the good performance of class-agnostic models in regions like Thalamus. Additional information on model evaluation, including class and structure specific performance, is given in Appendix 5. In particular, Supplementary Table 4 contains the sizes of these evaluation sets in each major structure, and Supplementary Section 7 contains the structure- and class specific losses.

**Table 2:** 2a Losses from leave-one-out cross-validation of candidate models. **Bold** numbers are best for their major structure. 2b Empirical performance of selected EL model by data abundance. The model is more accurate in Cre-leaf combinations where it draws on more data. The dataset variable *𝒟* indicates the set of experiments used to model a given connectivity. For example, *I_c_*∩ *I_L_* means only experiments with a given Cre-line in a given leaf are used to model connectivity for the corresponding cell-class in that leaf, while *I_L_* means that all experiments in that leaf are used. *I_wt_* ∩ *I_M_* means all wild-type experiments in the major structure are used - this was the model in Knox et al. (2019).

| $\hat{f}$ | Mean Leaf-Cre<br>Mean | NW Major-Cre<br>NW | NW Leaf-Cre<br>NW | NW Leaf<br>NW | NW Major-WT<br>NW | NW Major<br>NW | EL |
| --- | --- | --- | --- | --- | --- | --- | --- |
| $\mathcal{D}$ | $I_c \cap I_L$ | $I_c \cap I_M$ | $I_c \cap I_L$ | $I_L$ | $I_{wt} \cap I_M$ | $I_M$ | $I_L$ |
| Isocortex | 0.239 | 0.252 | 0.234 | 0.279 | 0.274 | 0.274 | **0.228** |
| OLF | 0.193 | 0.233 | 0.191 | **0.135** | 0.179 | 0.179 | 0.138 |
| HPF | 0.175 | 0.332 | 0.170 | 0.205 | 0.228 | 0.228 | **0.153** |
| CTXsp | **0.621** | **0.621** | **0.621** | **0.621** | **0.621** | **0.621** | **0.621** |
| STR | 0.131 | **0.121** | 0.128 | 0.169 | 0.232 | 0.232 | 0.124 |
| PAL | 0.203 | 0.205 | 0.203 | 0.295 | 0.291 | 0.291 | **0.188** |
| TH | 0.673 | 0.664 | 0.673 | **0.358** | 0.379 | 0.379 | 0.369 |
| HY | 0.360 | 0.382 | 0.353 | 0.337 | 0.317 | 0.317 | **0.311** |
| MB | 0.168 | 0.191 | 0.160 | 0.199 | 0.202 | 0.202 | **0.159** |
| P | 0.292 | 0.292 | 0.292 | 0.299 | 0.299 | 0.299 | **0.287** |
| MY | 0.268 | 0.347 | 0.268 | 0.190 | **0.189** | **0.189** | 0.204 |
| CB | **0.062** | **0.062** | **0.062** | 0.068 | 0.112 | 0.112 | 0.068 |

### Connectivities

We estimate matrices 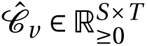 representing connections of source structures to target structures for particular Cre-lines *v* . We confirm the detection of several well-established connectivities within our tensor, although we expect additional interesting biological processes to be identifiable. The connectivity tensor and code to reproduce it are available at https://github.com/AllenInstitute/mouse_connectivity_models/tree/2020.

#### Overall connectivity

Several known biological projection patterns are evident in the wild-type connectivity matrix *𝒷_wt_* shown in Figure 2a. This matrix shows connectivity from leaf sources to leaf targets. Large-scale patterns like intraareal connectivities and ipsilateral connections between cortex and thalamus are clear, as in previous estimates in J. A. Harris et al. (2019); Knox et al. (2019); Oh et al. (2014). However, the layer-specific targeting of the different Cre-lines enables our estimated wild-type connectivities to display heterogeneity at the layer level. This contrasts the model in Knox et al. (2019), which is denominated as the NW Major-WT model whose accuracy is evaluated in Table 2a. For comparison, we also plot averages over component layers weighted by layer size projections between summary-structure sources and targets in the cortex in Figure 2b. Importantly, as shown in Table 2a this finer spatial resolution corresponds to the increased accuracy of our EL model over the NW Major-WT model.

**Figure 2:**
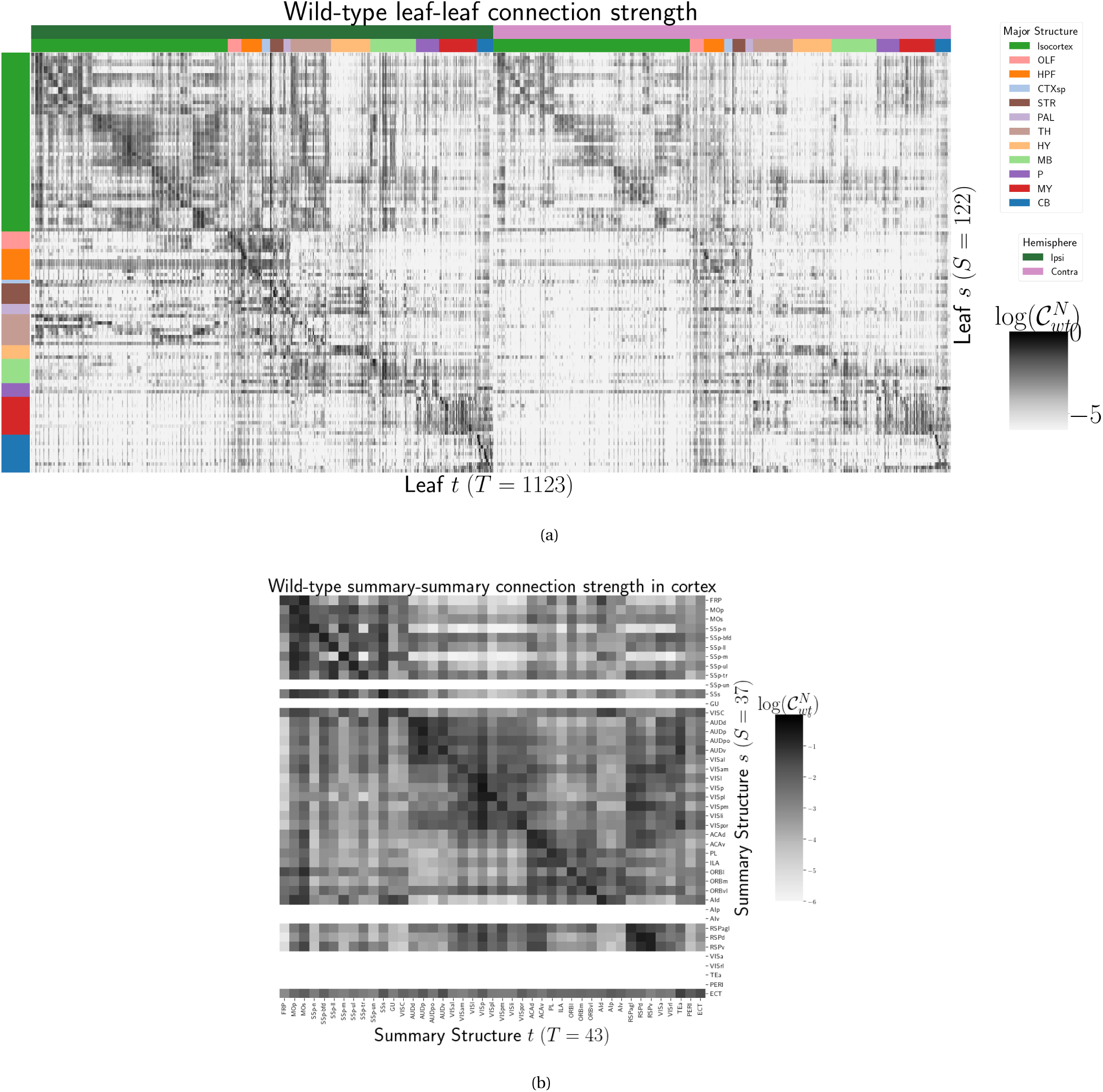
Wild-type connectivities. 2a Log wild-type leaf-to-leaf connectivity matrix log *𝒷* (*s*, *t*, *v_wt_*). Sources and target major brain division structure are shown. 2b Log wild-type intracortical connectivity matrix at the summary structure level. Summary structures without an injection centroid are left blank.

#### Class-specific connectivities

We investigate the presence of known biological processes within our connectivity estimates and confirm that these class-specific connectivities exhibit certain known behaviors. Although there is a rich anatomical literature using anterograde tracing data to describe projection patterns from subcortical sources to a small set of targets of interest, much of the accessible whole brain projection data is from the MCA project used here to generate the connectome models. Thus, we compare to external studies to validate our results while avoiding a circular validation of the data used to generate the model weights. The cell types and source areas with extensive previous anatomical descriptions of projections using both bulk tracer methods with cell type specificity and single cell reconstructions that we investigate are 1) thalamic-projecting neurons in the visual and motor cortical regions, 2) cholinergic neurons in the medial septum and nucleus of the diagonal band (MS/NDB); and 3) serotonergic neurons of the dorsal raphe nucleus (DR). Our estimated connections are in agreement with literature on these cell types.

##### Dependence of thalamic connection on cortical layer

Visual cortical areas VISp and VISl and cortical motor areas MOp and MOs have established layer-specific projection patterns that can be labeled with the layer-specific Cre-lines from the Allen datasets and others J. A. Harris et al. (2019); Jeong et al. (2016). Figure 3a shows that in VISp, the Ntsr1-Cre line strongly targets the core part of the thalamic LGd nucleus while in VISl, it a strong projection to the LP nucleus. In VISp, the Rbp4-Cre line strongly targets LP as well. Rbp4-Cre and Ntsr1-Cre injections target layers 5 and 6 respectively. Since we only generate connectivity estimates for structures with at least one injection centroid, this is shown by the position of non-zero rows in Figure 3a. To fill these gaps, as a heuristic alternative model, we also display an average connectivity matrix over all Cre-lines.

**Figure 3:**
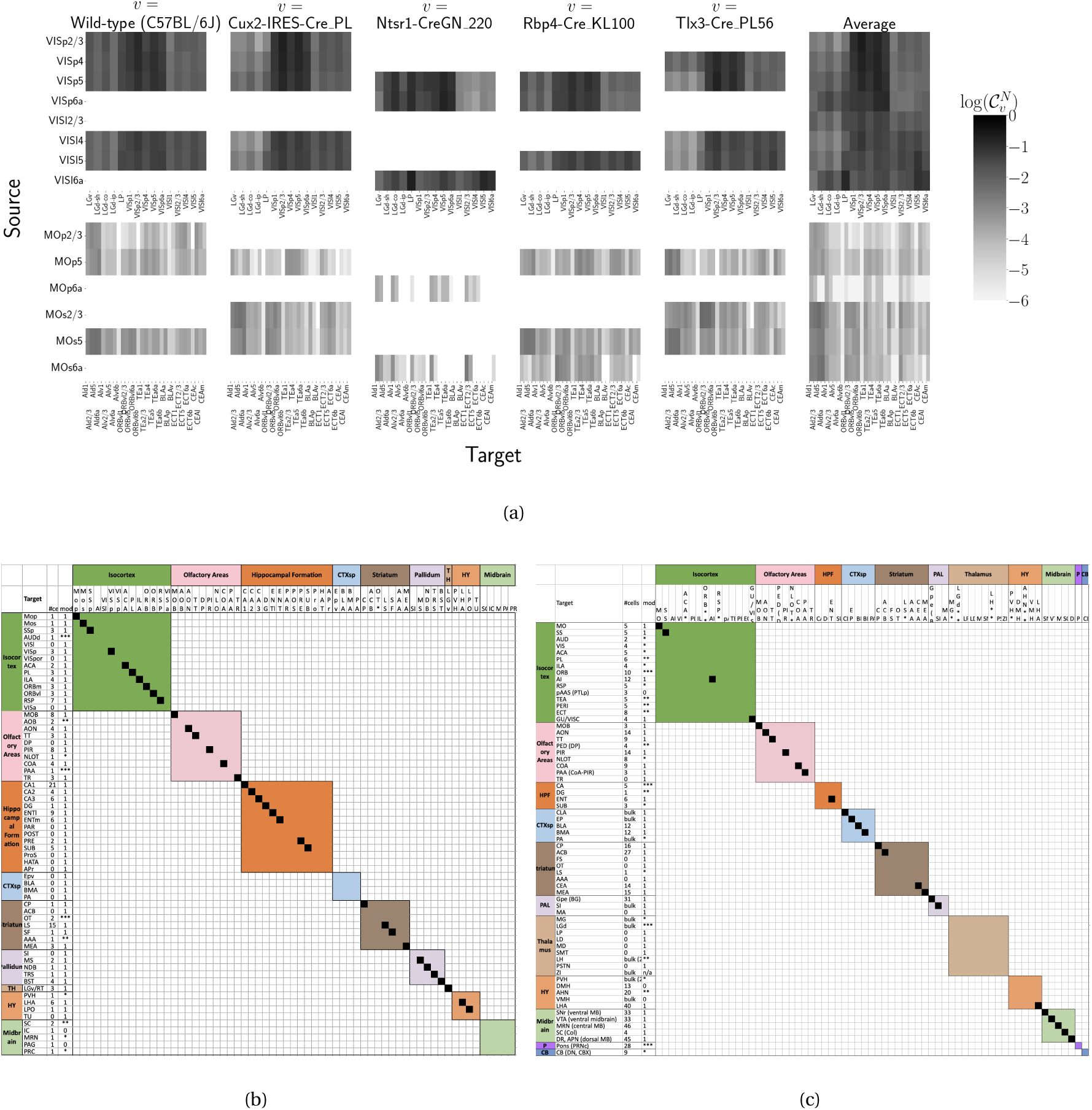
Cell-class specificity. 3a Selected cell-class and layer specific connectivities from two visual and two motor areas. Sources without an injection of a Cre-line are not estimated due to lack of data. 3b Targets reported for cholinergic cells in MS/NDB in Li et al. (2018) and above the 90th percentile in our Chat-IRES-Cre-neo model. A black box along the diagonal indicates the target was identified in both studies (*n =* 35). The number of single cells (out of 50) with a projection to each target is shown in the first column after the target acronym and the next column shows whether a target appeared in the 90% thresholded model weights (1= present, 0=absent). Some of these targets only appeared at lower thresholds as indicated by asterisks, *\*\* * >* 85*th*%, *\*\* >* 75*th*%, *\* >* 50*th*%. 3c Targets reported for serotonergic cells in DR in Ren et al. (2018, 2019) and above the 90th percentile in our Slc6a4-Cre_ET33 model.

##### MS and NDB projections in the Chat-IRES-Cre-neo model

Cholinergic neurons in the MS and NDB are well-known to strongly innervate the hippocampus, olfactory bulb, piriform cortex, entorhinal cortex, and lateral hypothalamus (Watson, Paxinos, & Puelles, 2012; Zaborszky et al., 2015). In the Allen MCA, cholinergic neurons were labeled by injections into Chat-IRES-Cre-neo mice. Figure 3b checks the estimated connectome weights to targets in these major brain divisions from MS and NDB. We observed that all these expected divisions were represented above the 90th percentile of weights from these source structures.

We also compared our Chat-IRES-Cre connectome model data for MS and NDB with the targets identified by Li et al. (2018). This single cell whole brain mapping project using Chat-Cre mice fully reconstructed n=50 cells to reveal these same major targets and also naming additional targets from MS/NDB (Li et al., 2018). We identified 150 targets at the fine leaf structure level among the top decile of estimated weights. To directly compare our data across studies, we merged structures as needed to get to the same ontology level, and remove ipsilateral and contralateral information. After formatting our data, we found 51 targets in the top 10%; Li et al. (2018) reported 47 targets across the 50 cells. There was good consistency overall between the target sets; 35 targets were shared, 12 were unique to the single cell dataset, and 16 unique to our model data. We checked whether targets missing from our dataset were because of the threshold level. Indeed, lowering the threshold to the 75th percentile confirmed 6 more targets-in-common, and all but 2 targets from Li et al. (2018) were above the 50th percentile weights in our model. Of note, the absence of a target in the single cell dataset that was identified in our model data is most likely due to the sparse sampling of all possible projections from only n=50 MS/NDB cells.

##### DR projections in the Slc6a4-Cre^.^ET33 model

Serotonergic projections from cells in the dorsal raphe (DR) are widely distributed and innervate many forebrain structures including isocortex and amygdala. In the Allen MCA, serotonergic neurons were labeled using Slc6a4-Cre_ET33 and Slc6a4-CreERT2_EZ13 mice. This small nucleus appears to contain a complex mix of molecularly distinct serotonergic neuron subtypes with some hints of subtype-specific projection patterns (Huang et al., 2019; Ren et al., 2018, 2019). We expect that the Cre-lines used here in the Allen MCA, which use the serotonin transporter promoter (Slc6a4-Cre and -CreERT2), will lead to expression of tracer in all the serotonergic subtypes recently described in an unbiased way, but this assumption has not been tested directly. We compared our model data to a single cell reconstruction dataset consisting of n=50 serotonergic cells with somas in the DR that also had bulk tracer validation 3c. After processing our data to match the target structure ontology level across studies, we identified 37 targets from the DR with weights above the 90th percentile, whereas Ren et al. (2019) listed 55 targets across the single cell reconstructions. Twenty seven of these targets where shared.

Overall there was good consistency between targets in olfactory areas, cortical subplate, CP, ACB and amygdala areas, as well in palidum and midbrain, while the two major brain divisions with the least number of matches are the isocortex and thalamus. There are a few likely reasons for these observations. First, in the isocortex, there is known to be significant variation in the density of projections across different locations, with the strongest innervation in lateral and frontal orbital cortices Ren et al. (2019). Indeed, when we lower the threshold and check for weights of the targets outside of the 90%, we see all but one of these regions (PTLp, parietal cortex which is not frontal or lateral) has a weight assigned in the top half of all targets. In the thalamus, our model predicted strong connections to several medial thalamic nuclei (i.e., MD, SMT) that were not targeted by the single cells. This discrepancy may be at least partially explained by the complex topographical organization of the DR that, like the molecular subtypes, is not yet completely understood. A previous bulk tracer study that specifically targeted injections to the central, lateral wings, and dorsal subregions of the DR reported semi-quantitative differences in projection patterns (Muzerelle, Scotto-Lomassese, Bernard, Soiza-Reilly, & Gaspar, 2016).Notably, Muzerelle et al. (2016) report that cells in the ventral region of DR project more strongly to medial thalamic nuclei, whereas the lateral and dorsal DR cells innervate more lateral regions (e.g., LGd). Thus, it is possible that the single cell somas did not adequately sample the entire DR.

### Connectivity Analyses

While the manual analysis in Section 3 is valuable for validation, scaling our interpretation of our connectivity estimates motivated us to apply dimension reduction methods to our connectivity estimates and understand whether the learned structure agrees with the expected biology. For example, Supplemental Section 7 shows a collection of connectivity strengths generated using Cre-specific models for wild-type, Cux2, Ntsr1, Rbp4, and Tlx3 cre-lines from VIS areas at leaf level in the cortex to cortical and thalamic nuclei. Sorting source and target structure/cell-class combinations hierarchical clustering shows, for example, that layer 6-specific Ntsr1 Cre-lines source regions cluster together. This makes sense, since layer 6-specific Ntsr1 Cre-line distinctly projects to thalamic nuclei, regardless of source summary structure, and in contrast with the tendency of other cell-classes to project to nearby regions within the cortex.

In contrast with heirarchical clustering, non-negative matrix factorization provides a simpler linear-model based factorization (Hastie, Tibshirani, & Friedman, n.d.). The low-dimensional coordinates returned by NMF nevertheless highlight important features within the connectivity matrix in a data-driven way. The learned projection archetypes *H* and model weights *W* are plotted in Figure 4. Intrastructural connections such as MB-MB MY-MY are visible in the 7th and 11th archetypes, but there is no obvious layer-6 specific signal. These factors may be used to generate a reconstructed connectivity matrix using the implied statistical model. Despite the relatively small number of learned additive factors, comparing with Figure 2a shows that this reconstructed matrix has a relatively globally plausible structure. Supplemental Sections 6 and 7 contain quantitative performance of this model, and assessment of factorization stability to ensure the decomposition is reliable across computational replicates. These indicate that though the displayed coordinate are fewer than the optimum number, this optimum is still an order of magnitude smaller than the dimension of the matrix itself. The supplement also quantitatively shows differential association of projection archetypes in this model with projection vectors of sources from the Cux2, Ntsr1, Rbp4, and Tlx3 Cre-lines.

**Figure 4:**
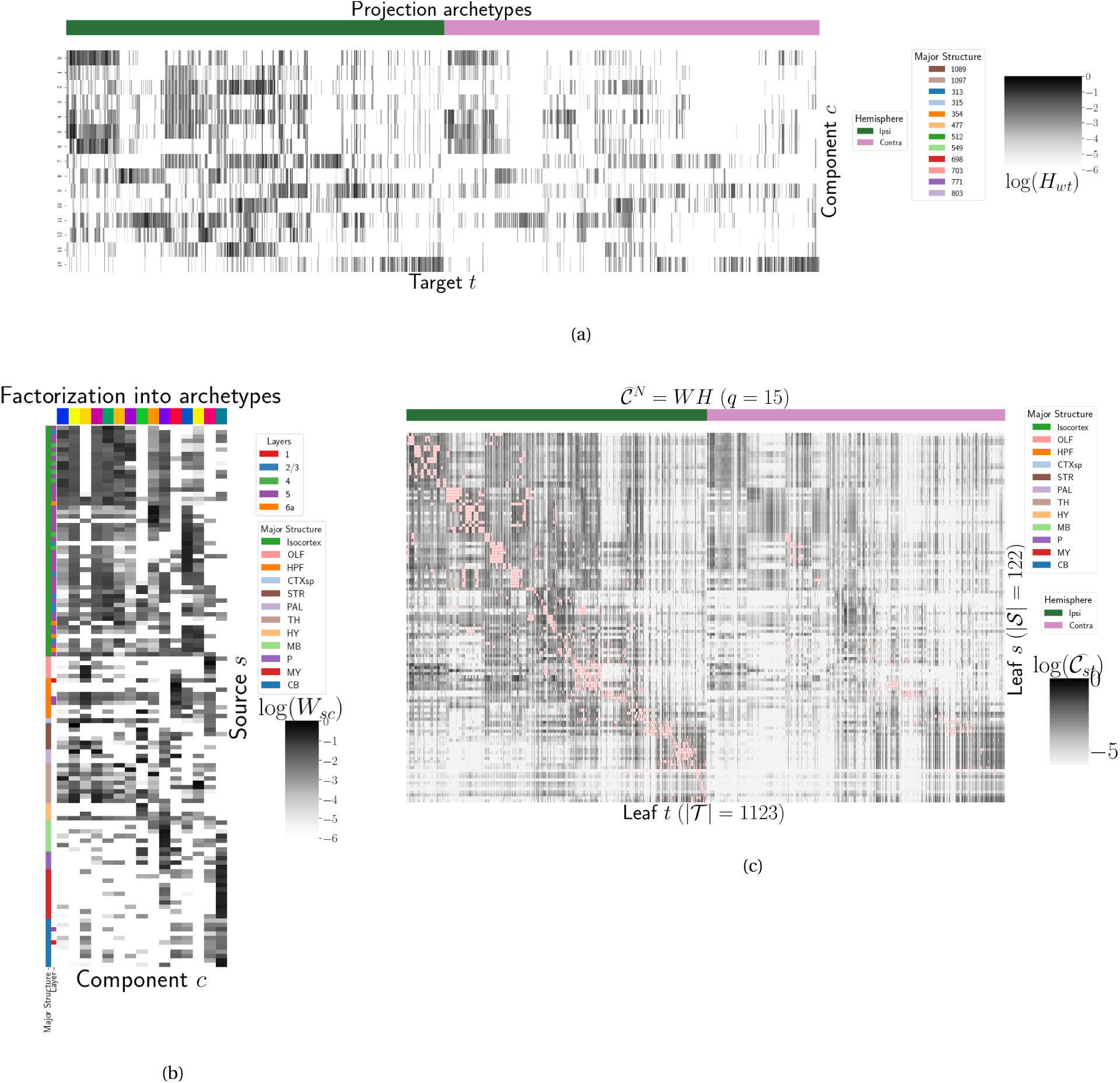
Non-negative matrix factorization results 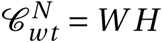 for *q =* 15 components. 4a Latent space coordinates *H* of *𝒷* . Target major structure and hemisphere are plotted. 4b Loading matrix *W* . Source major structure and layer are plotted. 4c Reconstruction of the normalized distal connectivity strength using the top 15 archetypes. Areas less than 1500 *µm* apart are not modeled, and therefore shown in pink.

## 4 DISCUSSION

The model presented here is among the first cell-type specific whole brain projectome models for a mammalian species, and it opens the door for a large number of models linking brain structure to computational architectures. Overall, we find expected targets, based on our anatomical expertise and published reports, but underscore that the core utility of this bulk connectivity analysis is not only in validation of existing connection patterns, but also in identification of new ones. We note that although the concordance appeared stronger for the cholinergic cells than the serotonergic cells, any differences might still be explained by the lack of high quality “ground-truth” datasets to validate these Cre-line connectome models. It is important to note several limitations of the current analyses. Short-range connections can be affected by saturating signals near injection sites, as well as the segmentation algorithm capturing dendrites as well as axons. Furthermore, larger numbers of single cell reconstructions that saturate all possible projection types would be a better gold standard than the small number of cells reported here. Future iterations of connectome models may also take into account single cell axon projection data, or synthesize with retrograde tracing experiments.

The Nadaraya-Watson estimator using the cell-type space based on similarities of projections, and theoretical justification of the use of an intermediate shape-constrained estimator, provides an empirically useful new tool for categorical modeling. Ours is not the first cross-validation based model averaging method Gao, Zhang, Wang, and Zou (2016), but our use of shape-constrained estimator in target-encoded feature space is novel and fundamentally different from Nadaraya-Watson estimators that use an optimization method for selecting the weights (Saul & Roweis, 2003). The properties of this estimator and its relation to estimators fit using an optimization algorithm are therefore a possible future avenue of research (Groeneboom & Jongbloed, 2018; Salha & El Shekh Ahmed, n.d.). Since the impact of the virus depends on the particular injection region, a deep model such as Lotfollahi, Naghipourfar, Theis, and Alexander Wolf (2019) could be appropriate, provided enough data was available, but our sample size seems too low to utilize a fixed or mixed effect model generative model. In a sense both the non-negative least squares Oh et al. (2014) and NW models can be thought of as improvements over the structure-specific average, and so is also possible that a yet undeveloped residual-based data-driven blend of these models could provide improved performance. Finally, we note that a Wasserstein-based measure of injection similarity per structure could naturally combine both the physical simplicity of the centroid model while also incorporating the full distribution of the injection signal.

The factorization of the connectivity matrix could also be improved. Non-linear data transformations or matrix decompositions, or tensor factorizations that account for correlations between cell-types could better capture the true nature of latent neural connections (K. D. Harris et al., 2016). Cre- or layer-specific signal recovery as performed here could be used to evaluate a range of matrix decompositions. This could help for example to understand the influence of traveling fibers on the observed connectivity (Llano & Sherman, 2008). From a statistical perspective, stability-based method for establishing archetypal connectivities in NMF is similar to those applied to genomic data Kotliar et al. (2019); Wu et al. (2016). Regardless of statistical approach, as in genomics, latent low-dimensional organization in connectivity should inspire search for similarly parsimonious biological correlates.

## ACKNOWLEDGMENTS

We thank the Allen Institute for Brain Science founder, Paul G. Allen, for his vision, encouragement, and support.

This supplement is divided into information about our dataset, supplemental methods, and supplemental results. However, certain topics are revisited between sections. Thus, if a reader is interested in, say, non-negative matrix factorization, they may find relevant information in both methods and results.

## 5 SUPPLEMENTAL INFORMATION

Our supplementary information consists of abundances of leaf/Cre-line combinations, information about distances between structures, and the size of our restricted evaluation dataset.

### Cre/structure combinations in *𝒟*

This section describes the abundances of structure and Cre-line combinations in our dataset. That is, it indicates how many experiments in our dataset with a particular Cre-line have an injection centroid in a particular structure. Users of the connectivity matrices who are interested in a particular Cre-line or structure can see the quantity and type of data used to compute and evaluate that connectivity.

**Figure 5:**
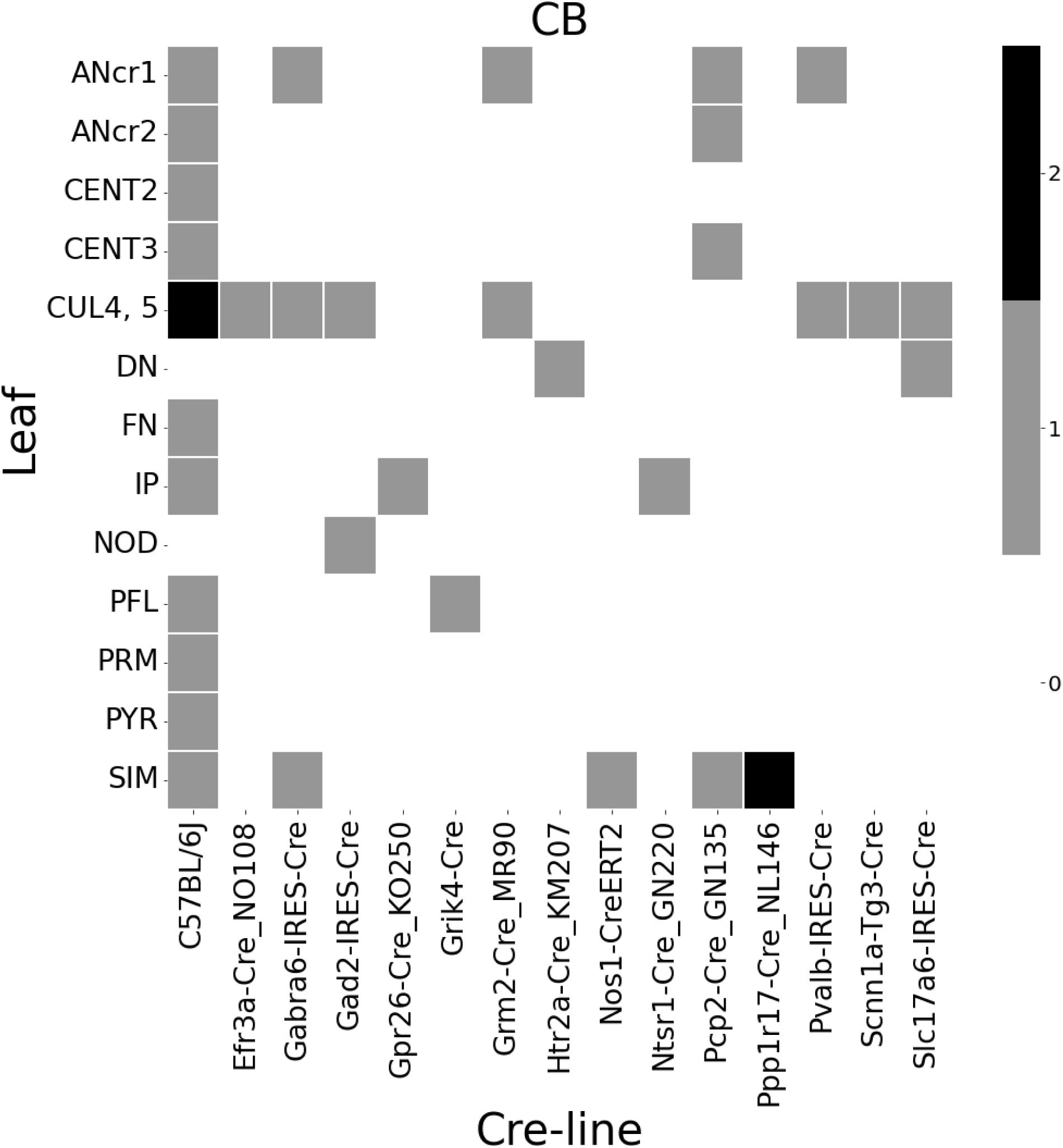
Frequencies of Cre-line and leaf-centroid combinations in our dataset.

**Figure 6:**
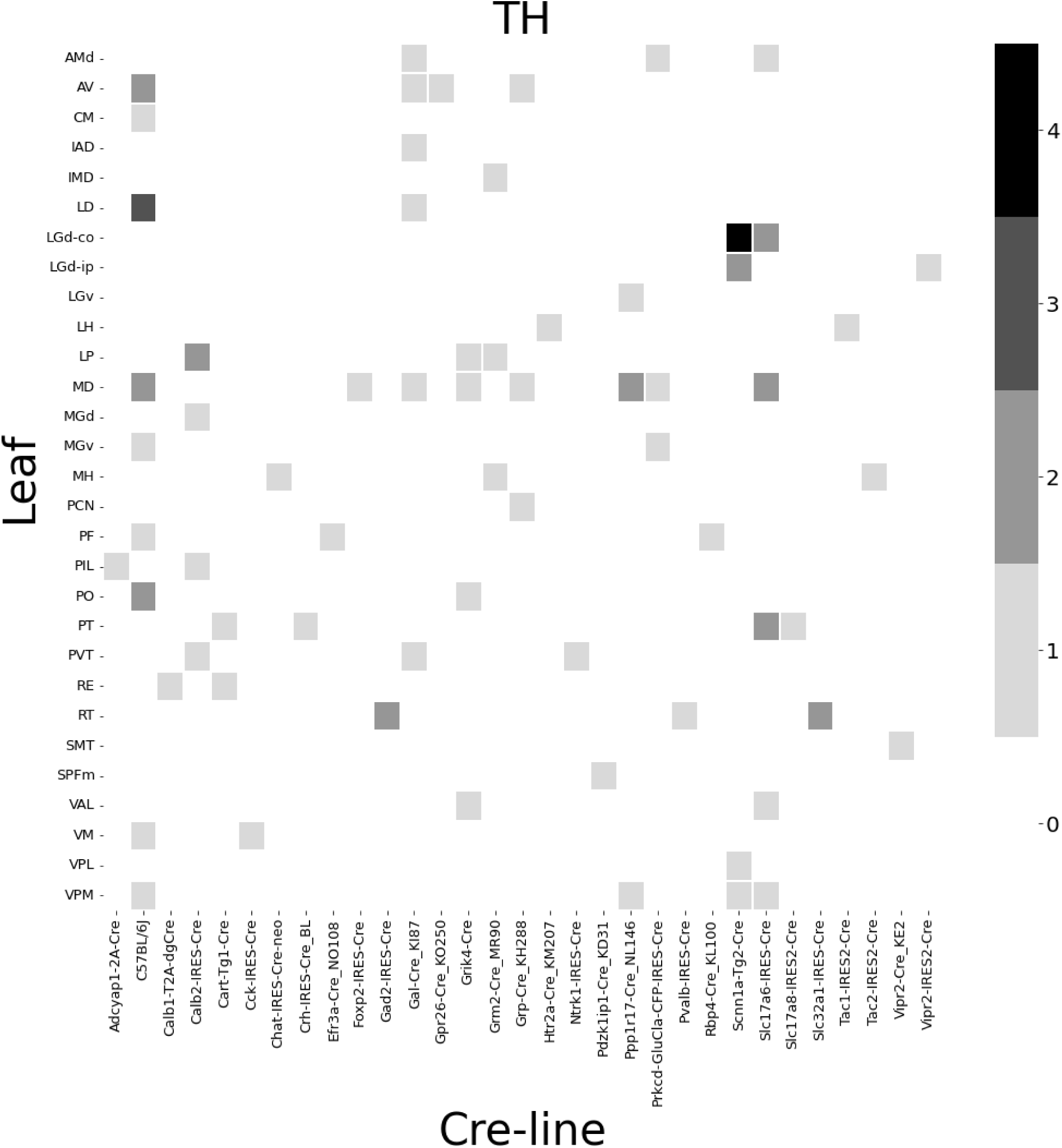
Frequencies of Cre-line and leaf-centroid combinations in our dataset.

**Figure 7:**
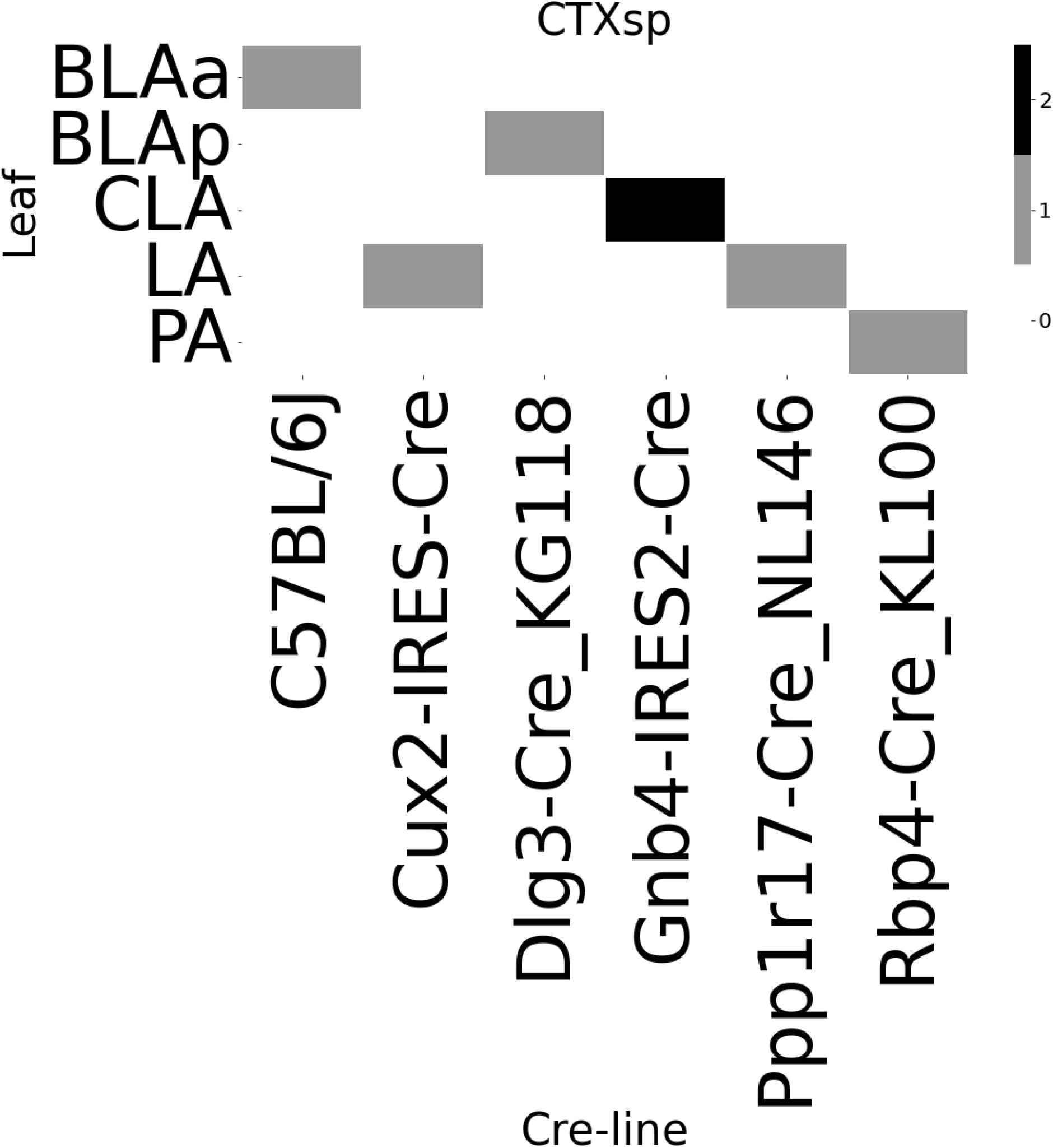
Frequencies of Cre-line and leaf-centroid combinations in our dataset.

**Figure 8:**
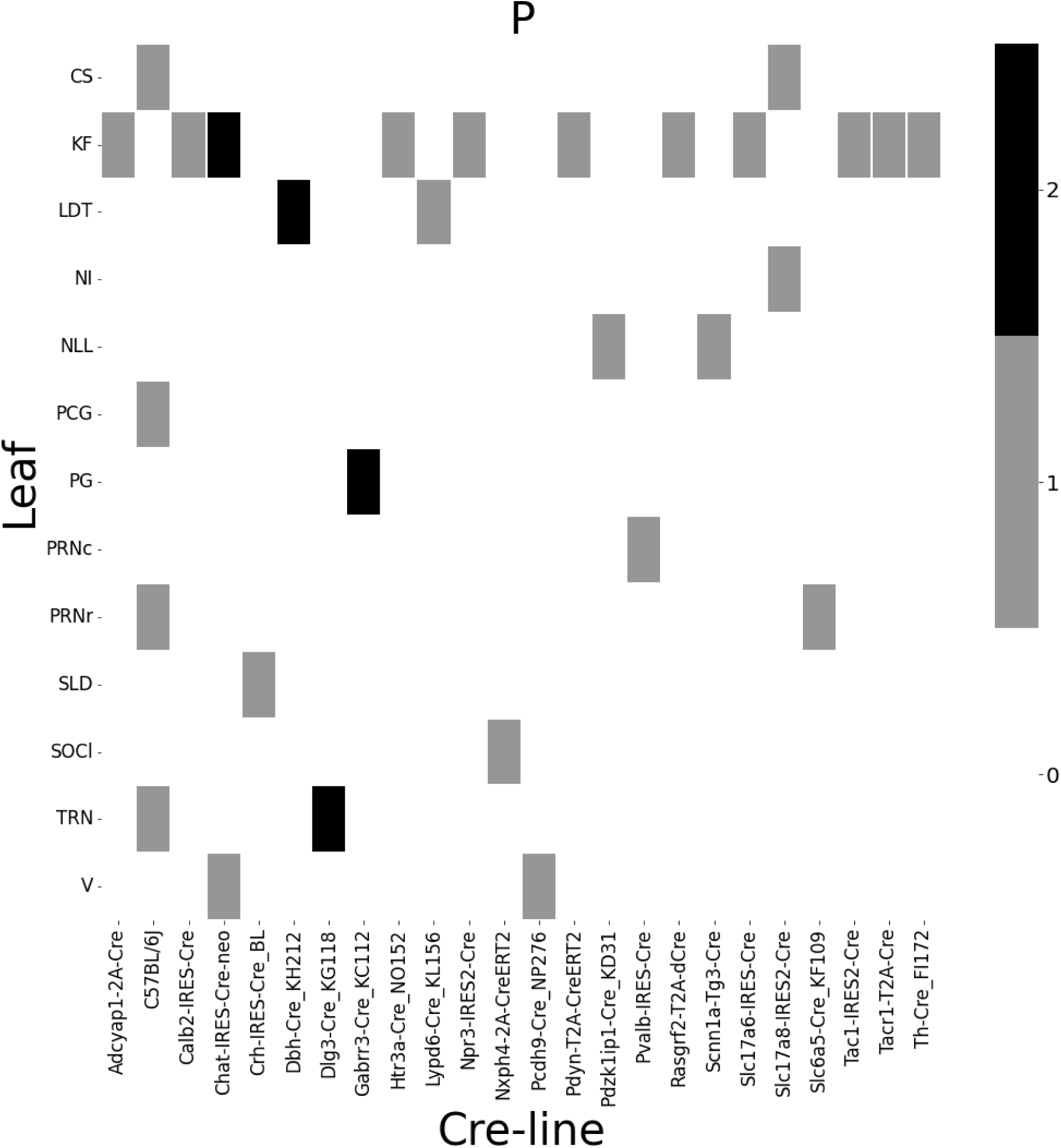
Frequencies of Cre-line and leaf-centroid combinations in our dataset.

**Figure 9:**
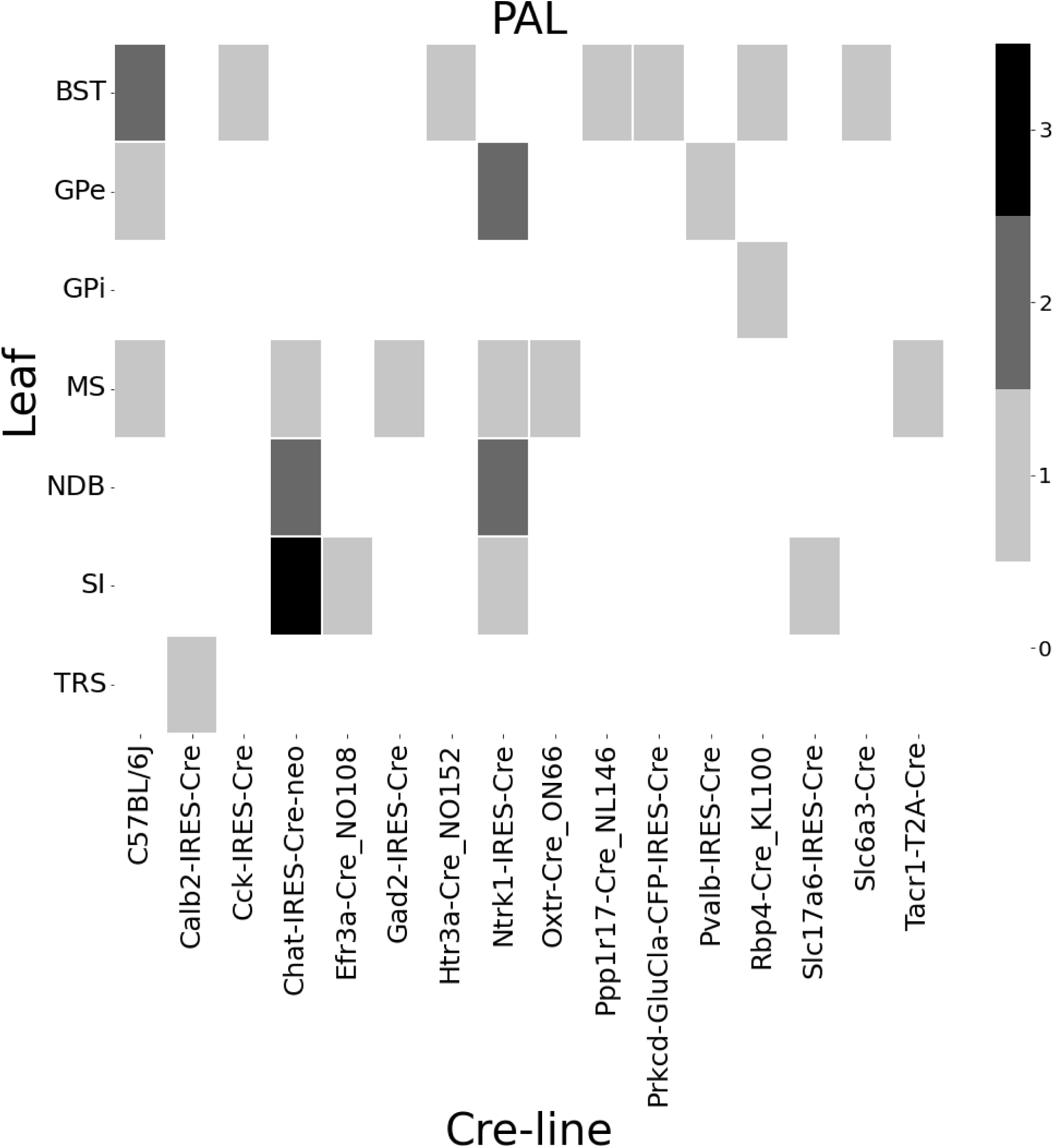
Frequencies of Cre-line and leaf-centroid combinations in our dataset.

**Figure 10:**
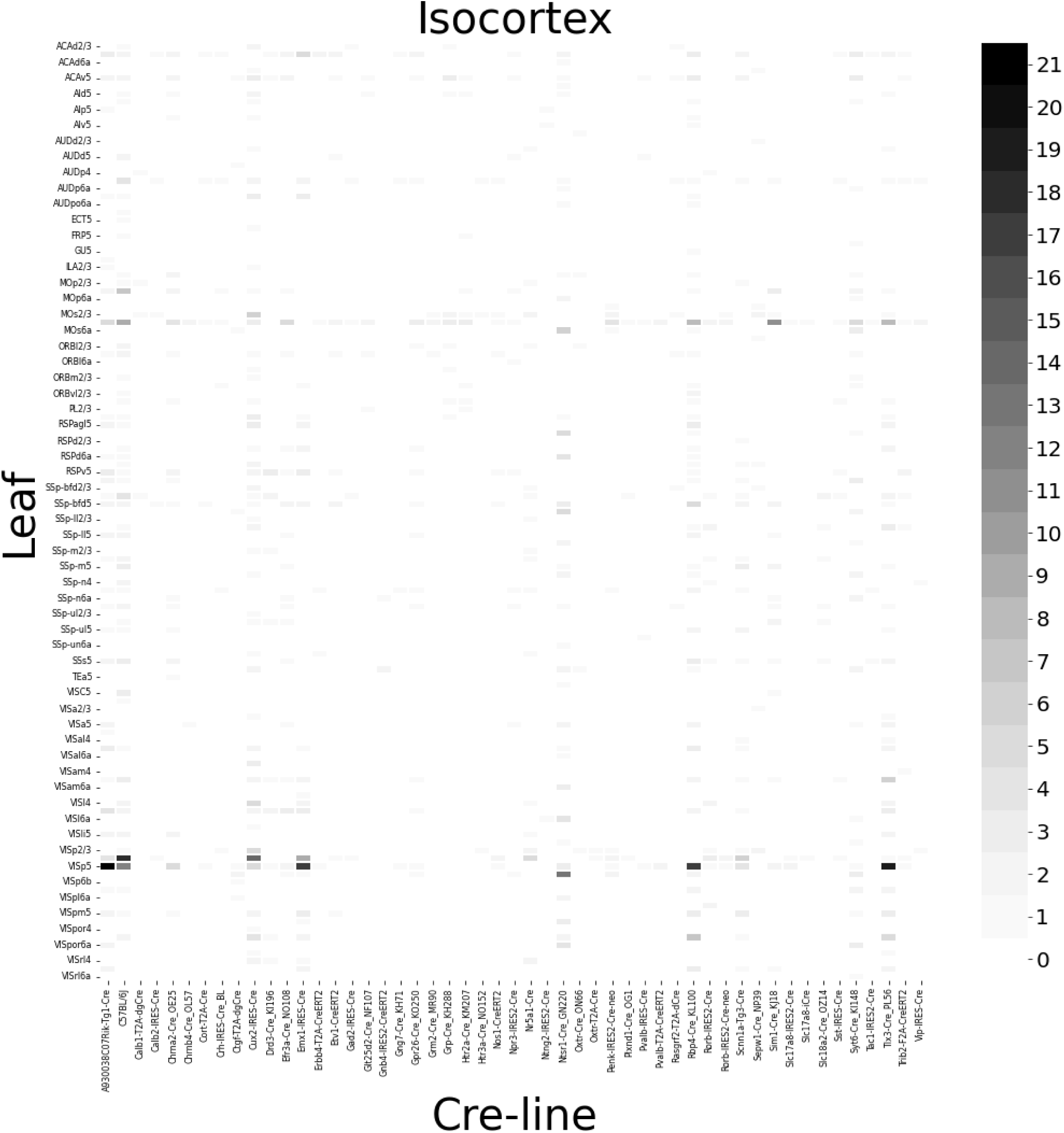
Frequencies of Cre-line and leaf-centroid combinations in our dataset.

**Figure 11:**
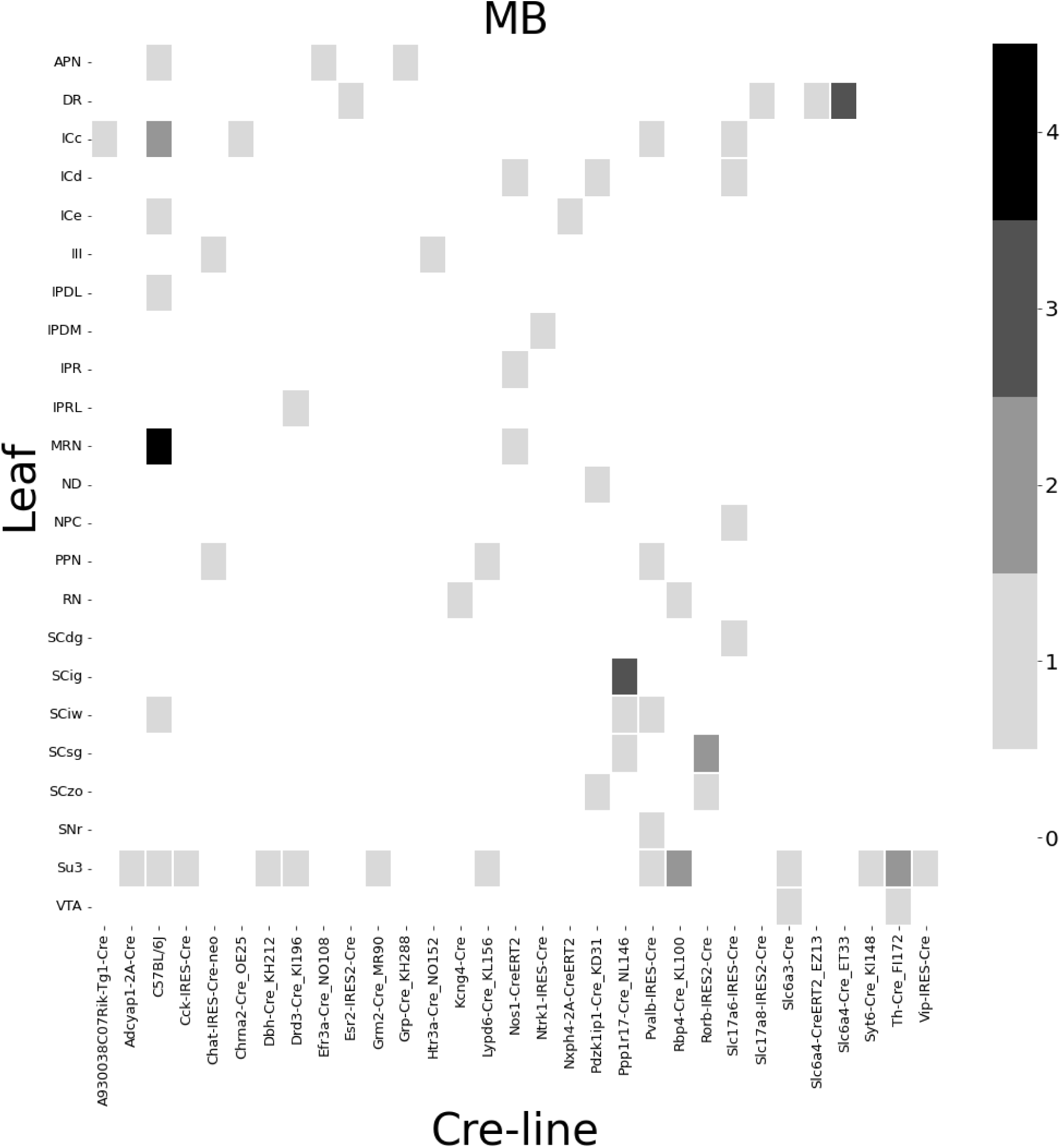
Frequencies of Cre-line and leaf-centroid combinations in our dataset.

**Figure 12:**
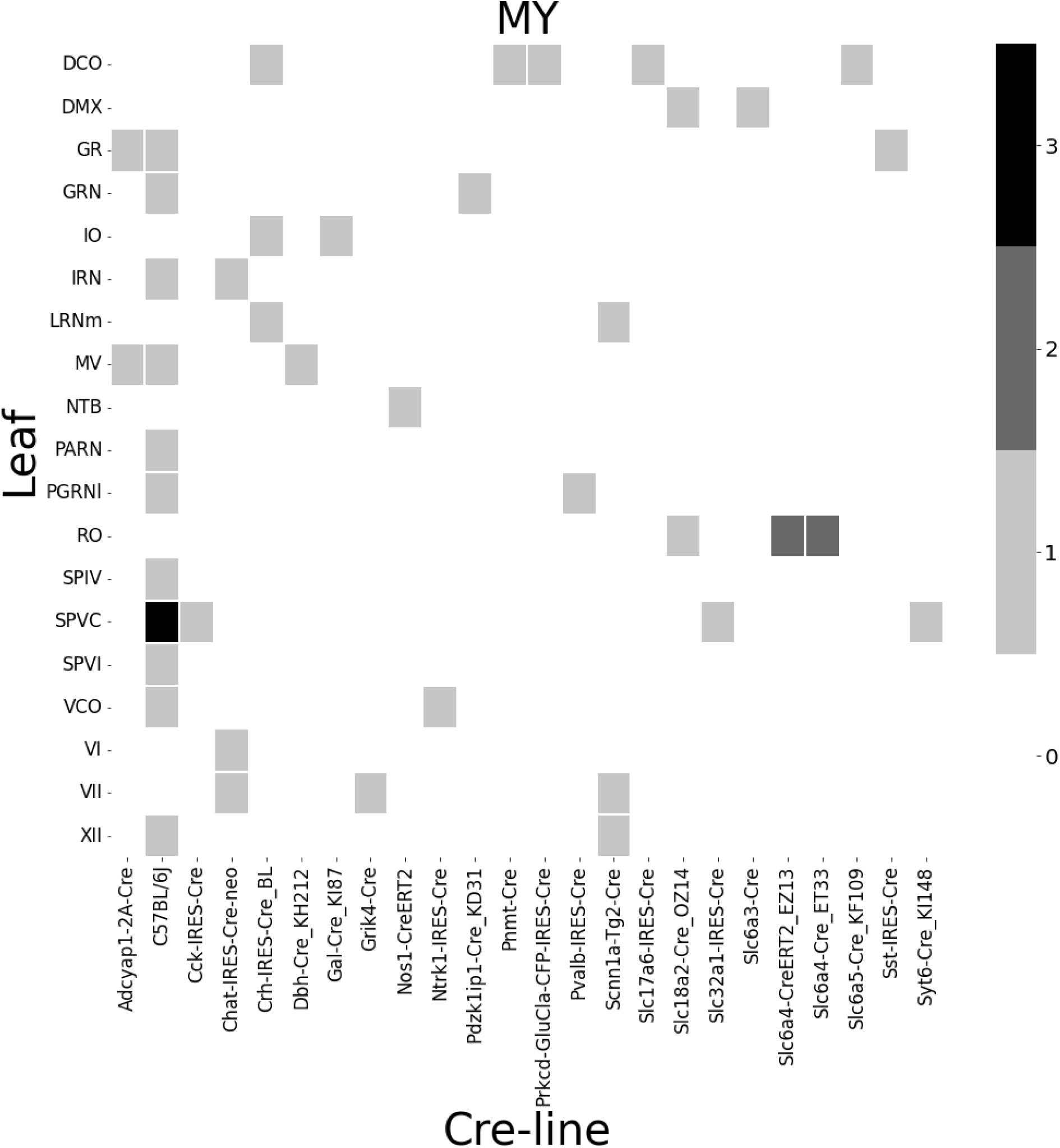
Frequencies of Cre-line and leaf-centroid combinations in our dataset.

**Figure 13:**
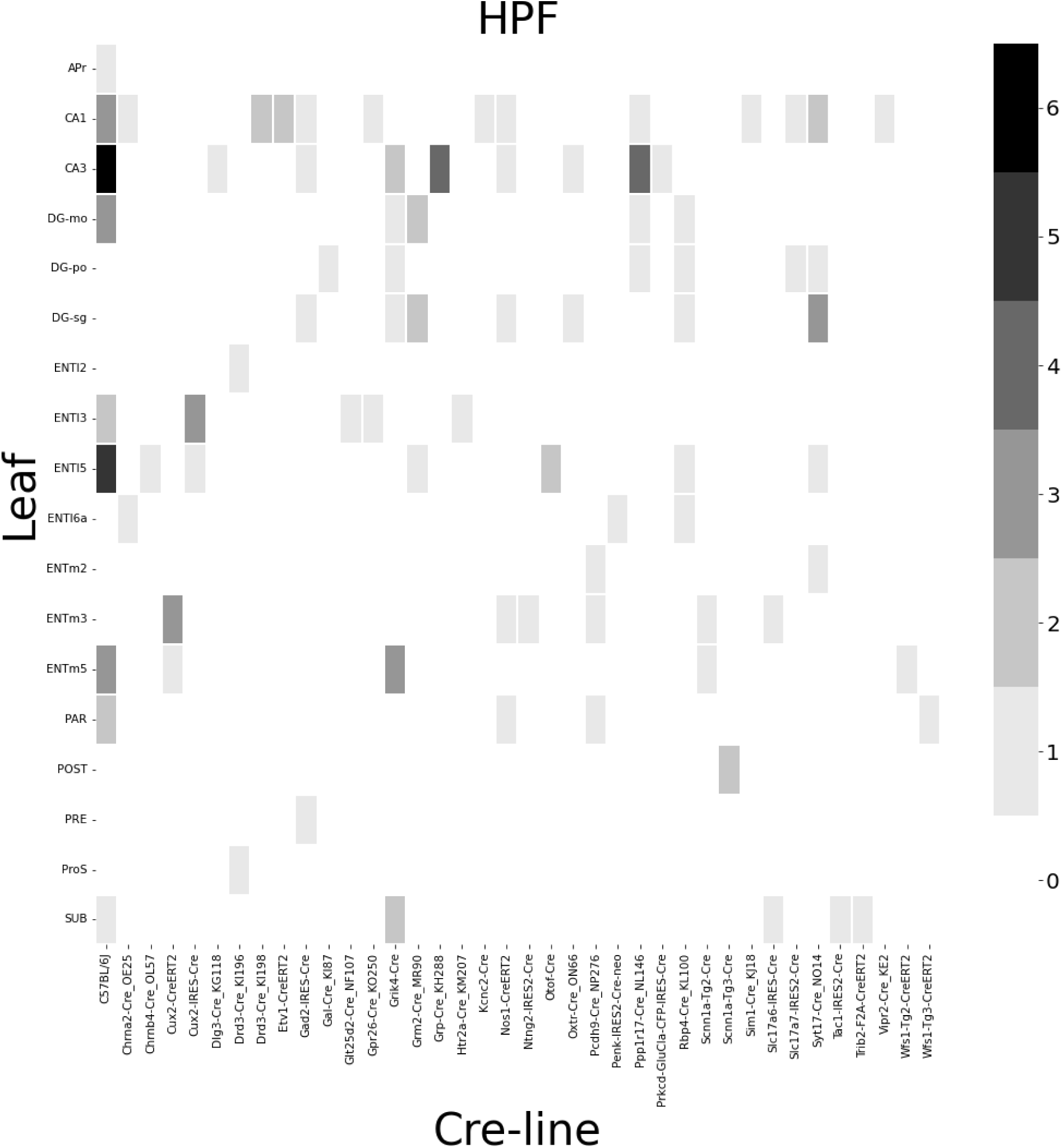
Frequencies of Cre-line and leaf-centroid combinations in our dataset.

**Figure 14:**
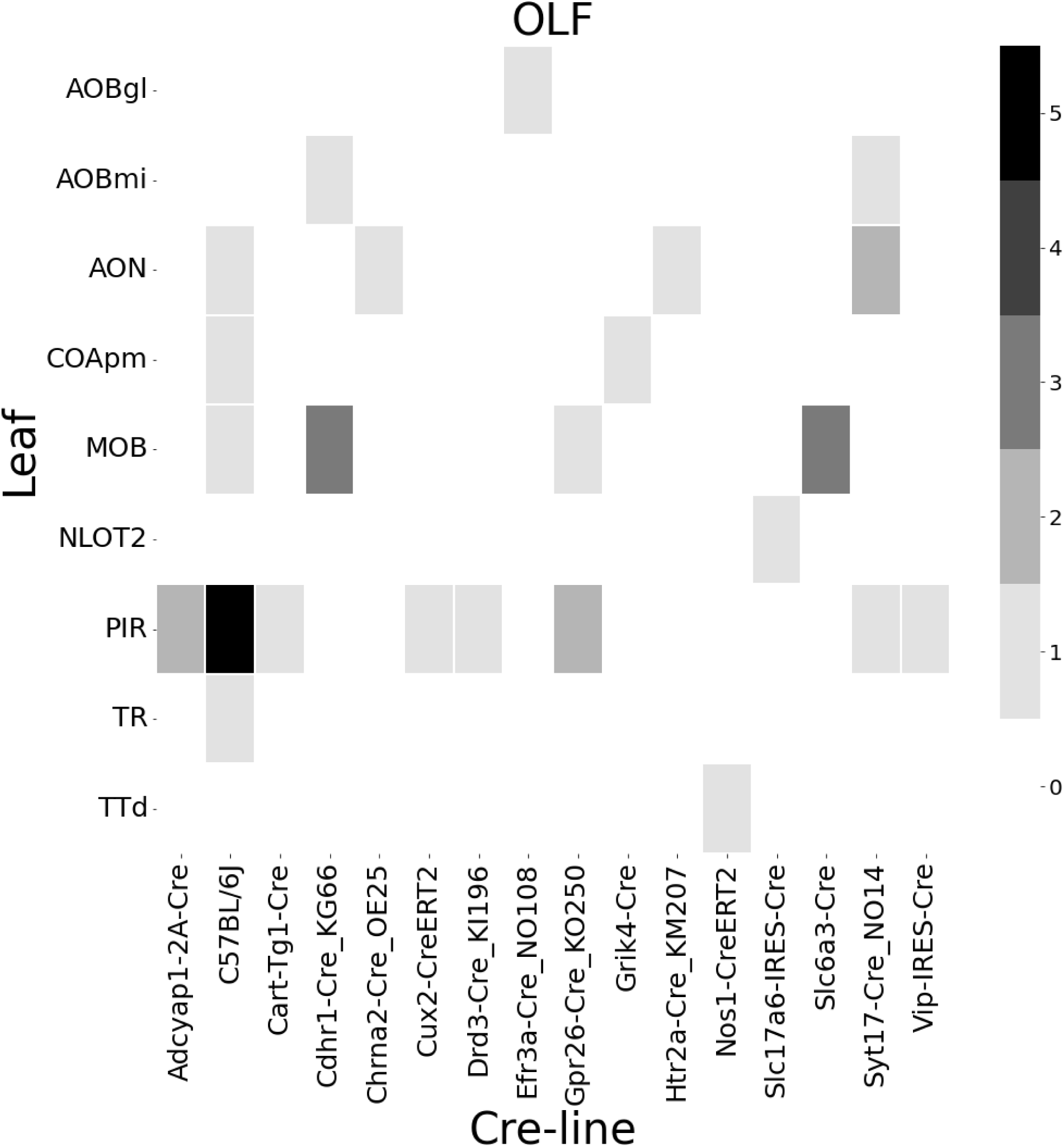
Frequencies of Cre-line and leaf-centroid combinations in our dataset.

**Figure 15:**
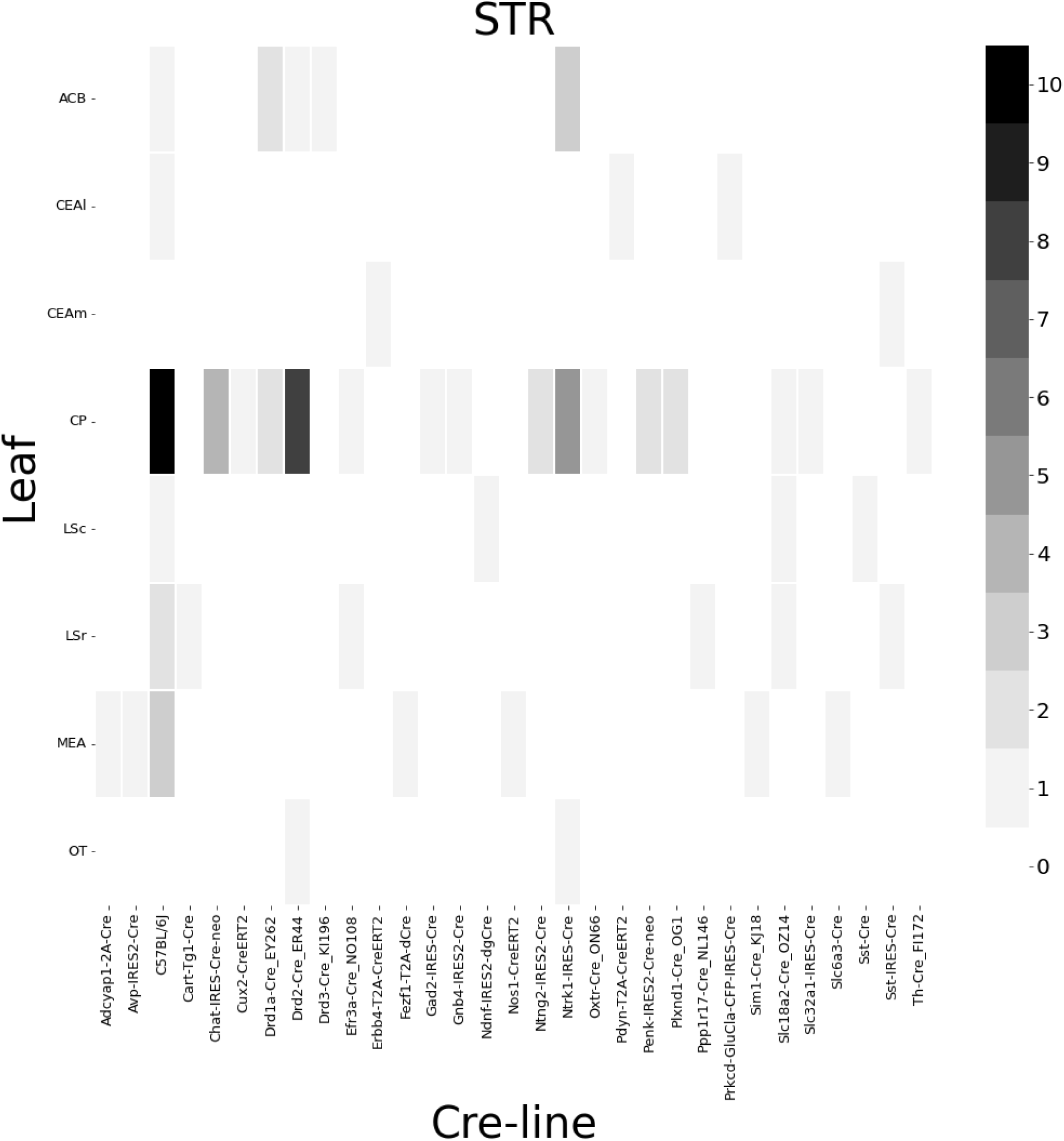
Frequencies of Cre-line and leaf-centroid combinations in our dataset.

**Figure 16:**
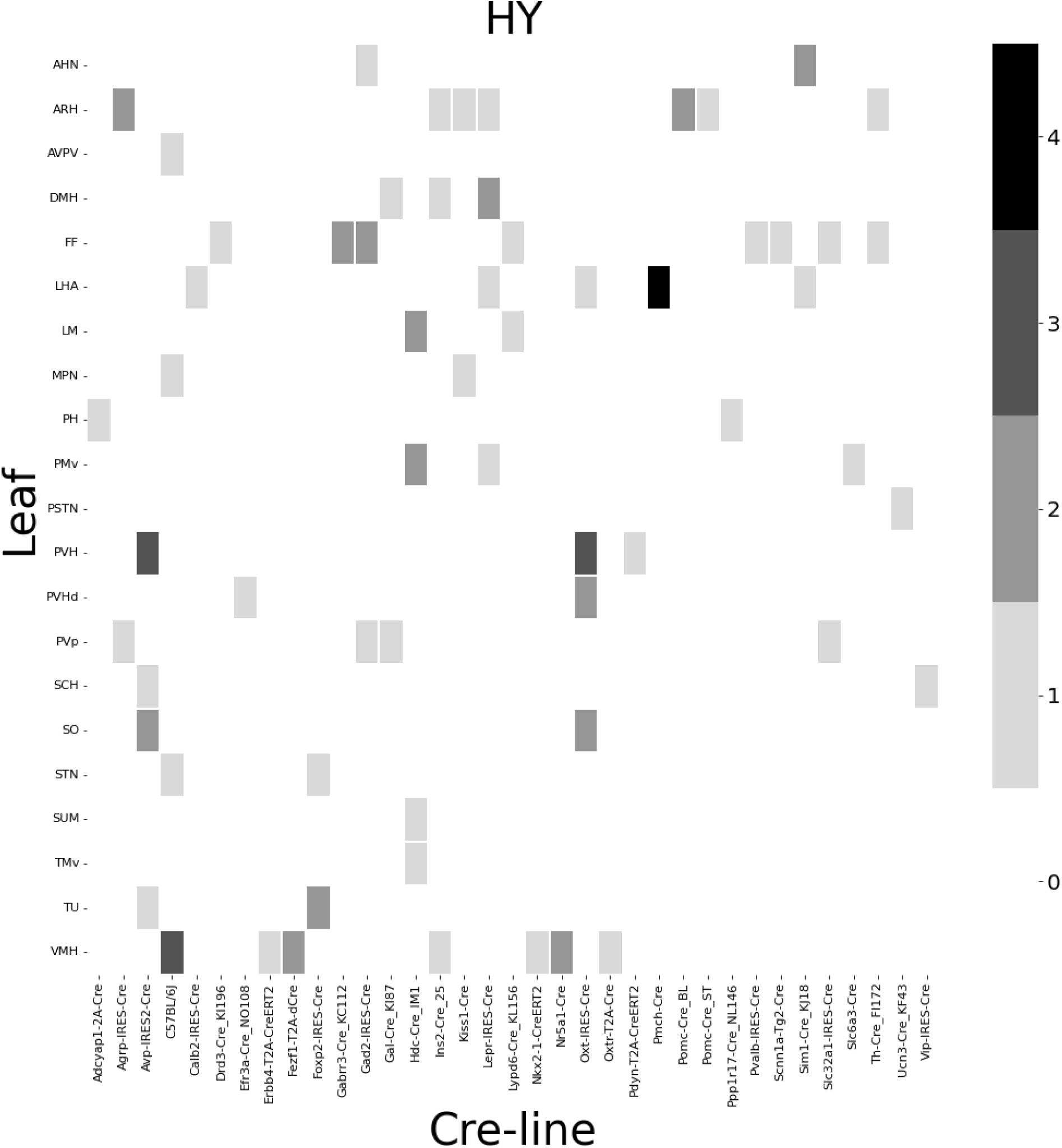
Frequencies of Cre-line and leaf-centroid combinations in our dataset.

### Distances between structures

The distance between structures has a strong effect on the connectivity (Knox et al., 2019). For reference, we show these distances here. Short range distances are not used in our matrix factorization approach. This masking is methodologically novel.

**Figure 17:**
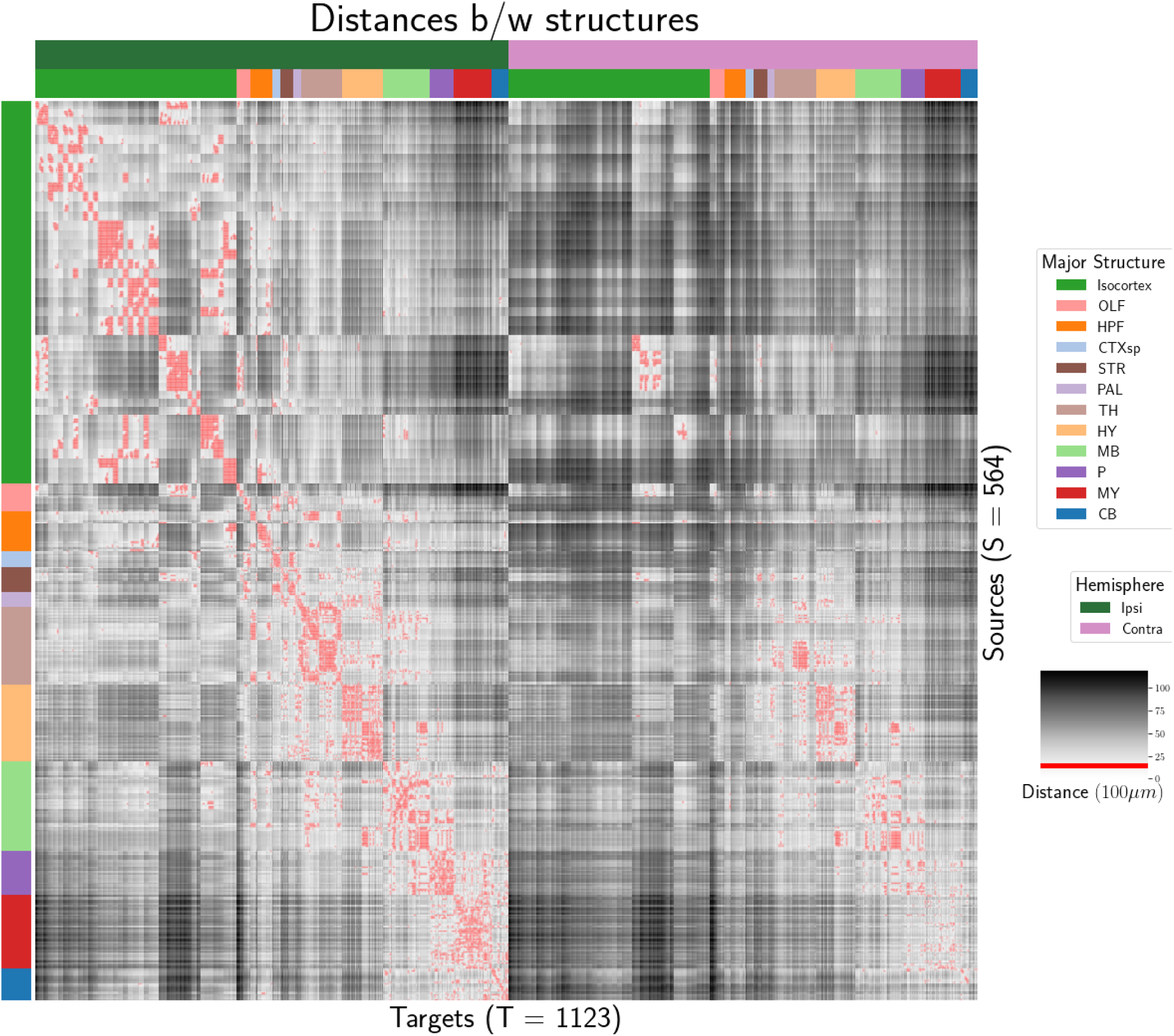
Distance between structures. Short-range connections are masked in red.

### Model evaluation

We give the sizes of our evaluation sets in leave-one-out cross-validation and additional losses using the injection-based normalization scheme from Knox et al. (2019).

#### Number of experiments in evaluation sets

In order to compare between methods, we restrict to the smallest set of evaluation indices. That is, the set of experiments used to validate our models combinations that are those whose combination of Cre-line and injection centroid leaf are present at least twice, since one experiment at least must be held out. This means that our evaluation set is smaller in size than our overall list of experiments.

**Table 3:** Number of experiments available to evaluate models in leave-one-out cross validation. The size of the evaluation set is lower than the total number of experiments since models that rely on a finer granularity of modeling have less data available to validate with, and we restrict all models to the smallest evaluation set necessitated by any of the modes. In this case, the Expected Loss and Cre- NW models require at least two experiments to be present with a combination of injection centroid structure and Cre-line for leave-one-out cross-validation. We also include results with a slightly smaller evaluation set that removes experiments without a sufficiently strong injection signal, as in Knox et al. (2019)

|  | Total | Evaluation set | Injection-thresholded evaluation |
| --- | --- | --- | --- |
| Isocortex | 1128 | 732 | 709 |
| OLF | 35 | 17 | 17 |
| HPF | 122 | 62 | 59 |
| CTXsp | 7 | 2 | 2 |
| STR | 78 | 45 | 44 |
| PAL | 30 | 11 | 11 |
| TH | 83 | 29 | 29 |
| HY | 85 | 41 | 38 |
| MB | 68 | 18 | 18 |
| P | 33 | 8 | 8 |
| MY | 46 | 7 | 7 |
| CB | 36 | 4 | 4 |

#### Injection-normalized losses

To compare with the injection-normalization procedure from Knox et al. (2019), we also remove experiments with small injection, and here give results for this slightly reduced set using injection-normalization. That is, instead of dividing the projection signal of each experiment by its *l* 1 norm (as we have used throughout the study), we divide by the *l* 1 norm of the corresponding injection signal. We find that setting a summed injection-signal of threshold of 1 is sufficient for evading pathological edge cases in this normalization, while still retaining a large evaluation set.

**Table 4:** Losses from leave-one-out cross-validation of candidate for injection-normalized regionalized connectivity on injection-thresholded evaluation set. **Bold** numbers are best for their major structure.

| $\hat{f}$ | Mean | NW | | | | | EL |
| --- | --- | --- | --- | --- | --- | --- | --- |
| $\mathcal{D}$ | $I_c \cap I_L$ | $I_c \cap I_M$ | $I_c \cap I_L$ | $I_L$ | $I_{wt} \cap I_M$ | $I_M$ | $I_L$ |
| Isocortex | 0.413 | 0.453 | 0.408 | 0.538 | 0.528 | 0.528 | **0.396** |
| OLF | 0.499 | 0.504 | 0.494 | 0.441 | 0.543 | 0.543 | **0.437** |
| HPF | 0.336 | 0.483 | 0.332 | 0.444 | 0.501 | 0.501 | **0.321** |
| CTXsp | 0.497 | 0.497 | 0.497 | 0.497 | 0.497 | 0.497 | 0.497 |
| STR | 0.359 | 0.386 | 0.359 | 0.364 | 0.433 | 0.433 | **0.322** |
| PAL | 0.519 | 0.497 | 0.519 | 0.436 | 0.459 | 0.459 | **0.434** |
| TH | 0.769 | 0.767 | 0.769 | **0.514** | 0.539 | 0.539 | 0.556 |
| HY | 0.414 | 0.439 | 0.414 | 0.441 | 0.452 | 0.452 | **0.399** |
| MB | 0.459 | 0.396 | 0.397 | 0.358 | **0.324** | **0.324** | 0.403 |
| P | **0.562** | **0.562** | **0.562** | 0.758 | 0.764 | 0.764 | **0.562** |
| MY | 0.699 | 0.552 | 0.621 | **0.439** | 0.578 | 0.578 | **0.439** |
| CB | 0.849 | 0.689 | 0.849 | 0.500 | 0.615 | 0.615 | **0.495** |

#### Projection-normalized losses on thresholded set

We also give results for the projection-normalization procedure from the main text on this reduced subset.

**Table 5:** Losses from leave-one-out cross-validation of candidate for normalized regionalized connectivity on injection-thresholded evaluation set. **Bold** numbers are best for their major structure.

| $\hat{f}$ | Mean | NW | | | | | EL |
| --- | --- | --- | --- | --- | --- | --- | --- |
| $\mathcal{D}$ | $I_c \cap I_L$ | $I_c \cap I_M$ | $I_c \cap I_L$ | $I_L$ | $I_{wt} \cap I_M$ | $I_M$ | $I_L$ |
| Isocortex | 0.229 | 0.248 | 0.224 | 0.274 | 0.269 | 0.269 | **0.217** |
| OLF | 0.193 | 0.233 | 0.191 | **0.135** | 0.179 | 0.179 | 0.138 |
| HPF | 0.178 | 0.342 | **0.172** | 0.212 | 0.235 | 0.235 | **0.172** |
| CTXsp | **0.621** | **0.621** | **0.621** | **0.621** | **0.621** | **0.621** | **0.621** |
| STR | 0.128 | **0.117** | 0.124 | 0.171 | 0.234 | 0.234 | 0.125 |
| PAL | 0.203 | 0.205 | 0.203 | 0.295 | 0.291 | 0.291 | **0.188** |
| TH | 0.673 | 0.664 | 0.673 | **0.358** | 0.379 | 0.379 | 0.417 |
| HY | 0.358 | 0.378 | 0.351 | 0.331 | 0.312 | **0.312** | 0.314 |
| MB | 0.168 | 0.191 | **0.160** | 0.199 | 0.202 | 0.202 | **0.160** |
| P | 0.292 | 0.292 | 0.292 | 0.299 | 0.299 | 0.299 | **0.287** |
| MY | 0.268 | 0.347 | 0.268 | **0.167** | 0.189 | 0.189 | 0.196 |
| CB | 0.062 | 0.062 | 0.062 | 0.068 | 0.108 | 0.108 | **0.061** |

## 6 SUPPLEMENTAL METHODS

This section consists of additional information on preprocessing of the neural connectivity data, estimation of connectivity, and matrix factorization.

### Data preprocessing

Several data prepreprocessing steps take place prior to evaluations of the connectivity matrices. These steps are described in Algorithm PREPROCESS. The arguments of this normalization process - injection signals *x*(*i*), projection signals *y* (*i*), injection fraction *F* (*i*), and data quality mask *q*(*i*) - were downloaded using the Allen SDK, a programatic interface to the brain connectivity data. The injections and projection signals *ℬ →* [0, 1] were segmented manually in histological analysis. The projection signal gives the proportion of pixels within the voxel displaying fluorescence, and the injection signal gives the proportion of pixels within the histologically-selected injection subset displaying fluorescence. The injection fraction *F* (*i*): *ℬ →* [0, 1] gives the proportion of pixels within each voxel in the injection subset. Finally, the data quality mask *q*(*i*): *ℬ →*{0, 1} gives the voxels that have valid data.

Our preprocessing makes use of the above ingredients, as well as several other essential steps. First, we compute the weighted injection centroid

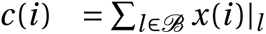

where *x*(*i*)*|_l_* is the injection density at location *l* ∈ **ℝ**^3^. Given a regionalization *ℛ* from the Allen SDK, we can also access regionalization map *ℛ* : *ℬ → **R***. This induces a functional of connectivities from the space of maps 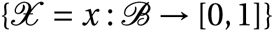

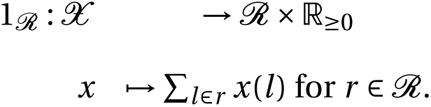

We also can restrict a signal to a individual structure as

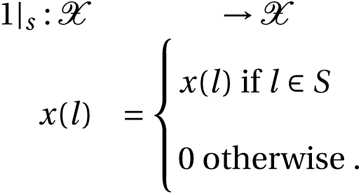

Finally, given a vector or array *a* ∈ **ℝ***^T^*, we have the *l* 1 normalization map

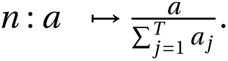

Denote *m* as the major structure containing an experiment, and define ʘ for maps *ℬ → **R***[0, 1] by e.g. (*y* (*i*) ʘ *q*(*i*))*|_l_* :*=* (*y* (*i*)*|_l_*)(*q*(*i*)*|_l_*). We then can write the preprocessing algorithm.

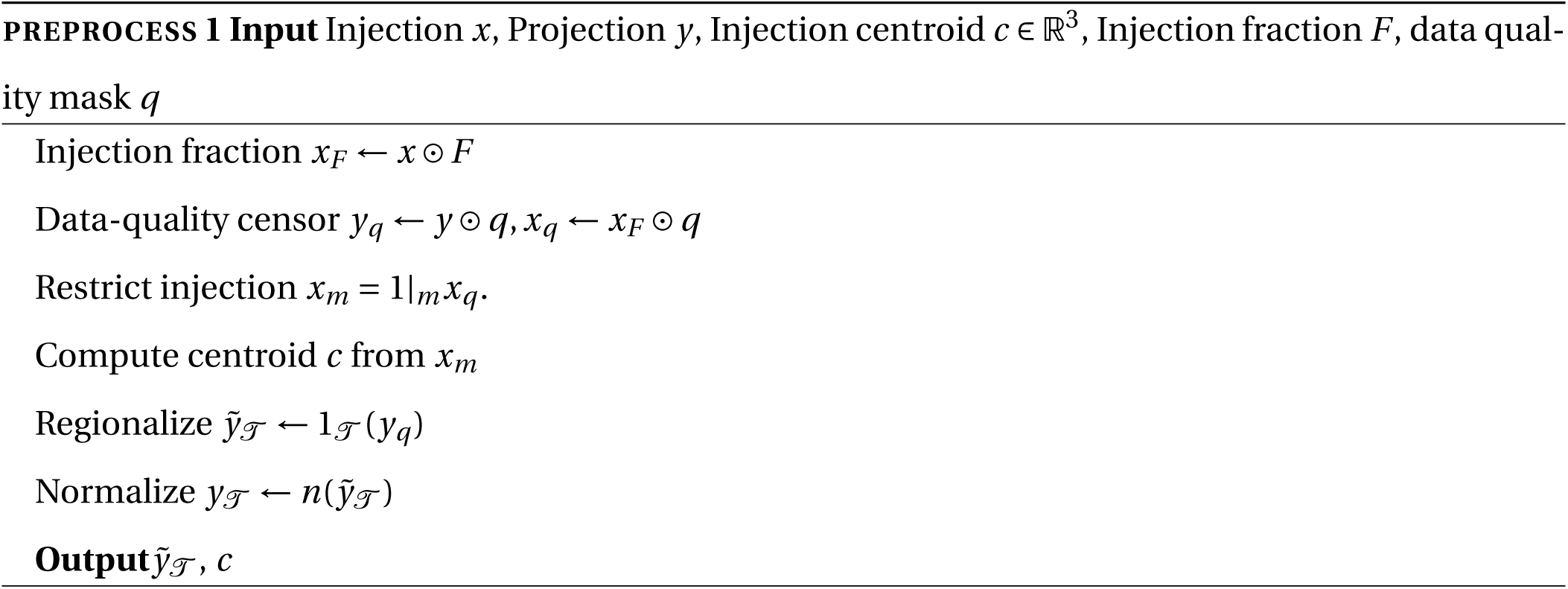

### Estimators

As mentioned previously, we can consider our estimators as modeling a connectivity vector 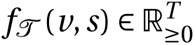. Thus, for the remainder of this section, we will discuss only *f* (*v*, *s*). We review the Nadaraya-Watson estimator from Knox et al. (2019), and describe its conversion into our cell-class specific Expected Loss estimator.

#### Centroid-based Nadaraya-Watson

In the Nadaraya-Watson approach of Knox et al. (2019), the injection is considered only through its centroid *c*(*i*), and the projection is considered regionalized. That is,

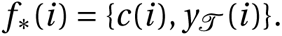

Since the injection is considered only by its centroid, this model only generates predictions for particular locations *l*, and the prediction for a structure *s* is given by integrating over locations within the structure

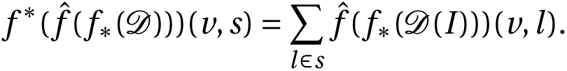

Here, *I* is the training data, and 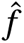 is the Nadaraya-Watson estimator

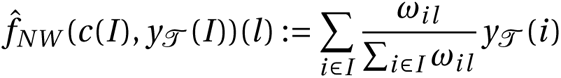

where *ω_il_* :*=* exp(−*γd* (*l*, *c*(*i*))^2^) and *𝒟* is the Euclidean distance between centroid *c*(*i*) and voxel with position *l*.

Several facets of the estimator are visible here. A smaller *γ* corresponds to a greater amount of smoothing, and the index set *I* ⊆ {1 : *n*} generally depends on *s* and *v* . Varying *γ* bridges between nearest neighbor prediction and averaging of all experiments in *I* . In Knox et al. (2019), *I* consisted of experiments sharing the same brain division, i.e. *I = I_m_*, while restricting of index set to only include experiments with the same cell class gives the class-specific Cre-NW model. Despite this restriction, we fit *γ* by leave-one-out cross-validation for each *m* rather than a smaller subset like *s* or *v* . That is,

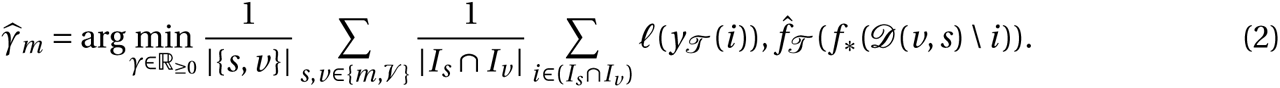

#### The Expected-Loss estimator

Besides location of the injection centroid, cell class also influences projection. Thus, we introduce method for estimating the effect of Cre-distance, which we define as the distance between the projections of the mean experiment of one (Cre,leaf) pair with another. Equivalently, relatively small Cre-distance defines what we call similar cell classes. This method assigns a predictive weight to each pair of training points that depends both on their centroid-distance and Cre-distance. This weight is determined by the expected prediction error of each of the two feature types

We define Cre-line behavior as the average regionalized projection of a Cre-line in a given structure (i.e. leaf). The vectorization of categorical information is known as **target encoding**

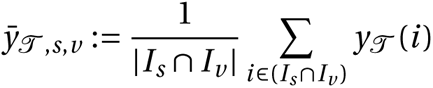

The Cre-distance is then define a **Cre-distance** in a leaf to be the distance between the target-encoded projections of two Cre-lines. The relative predictive accuracy of Cre-distance and centroid distance is determined by fitting a surface of projection distance as a function of Cre-distance and centroid distance. When we use shape-constrained B-splines to estimate this weight, the weights then may be said to be used in a Nadaraya-Watson estimator. For this reason, we call this the Expected Loss Estimator. The resulting weights are then utilized in a Nadaraya-Watson estimator in a final prediction step.

In mathematical terms, our full feature set consists of the centroid coordinates and the target-encoded means of the combinations of virus type and injection-centroid structure. That is,

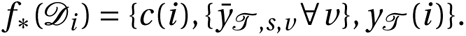

*f\** is defined as in (2). The expected loss estimator is then

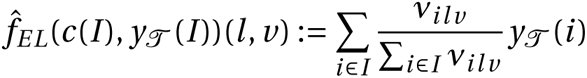

where

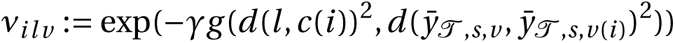

and *s* is the structure containing *l* .

The key step therefore is finding a suitable function *g* with which to weight the positional and (Cre,leaf) information. Note that *g* must be a concave, non-decreasing function of its arguments with with *g* (0, 0) *=* 0. Then, *g* defines a metric on the product of the metric spaces defined by experiment centroid and target-encoded cre-line, and 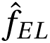 is a Nadaraya-Watson estimator. A derivation of this fact is given later in this section.

We therefore use a linear generalized additive model of shape-constrained B-splines to estimate *g* (Eilers & Marx, 1996). This is a method for generating a predictive model *g* that minimizes the loss of

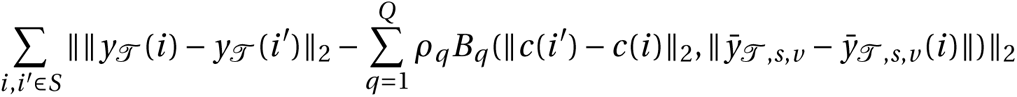

given the constraints on *g* . That is, given all pairs of experiments with injection centroid in the same structure, *g* gives a prediction of the distance between their projections made using the distance between the average behavior of their Cre-lines given their injection centroid, and the distance between their injection centroids. In particular, *g* is the empirically best such function within the class of *B* -splines, which Similarly to the Nadaraya-Watson model, we make the decision to fit a *g* separately for each major brain division, and select 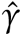 as in 2. We set *Q =* 10 and leave validation of this parameter, as well as the precise nature of the polynomial B-spline terms *B_q_* out of the scope of this paper. Empirically this leads to a smooth surface using the pyGAM Python package (Servén D., n.d.).

**Figure 18:**
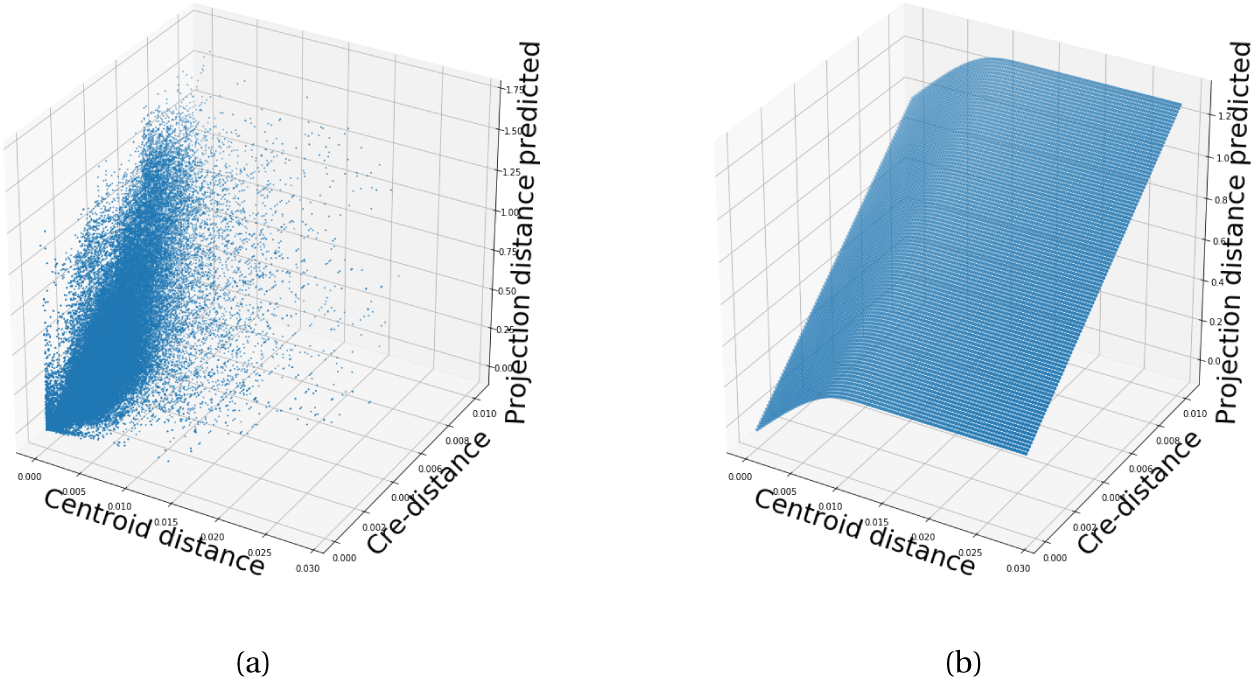
Fitting *g* . 18a Distribution of projection errors against centroid distance and cre-distance in Isocortex. 18b estimated 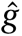 using B-splines. Projection distance is 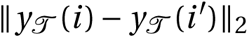, Cre-distance is 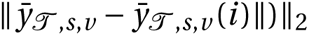, and centroid-distance is 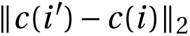.

#### Justification of shape constraint

The shape-constrained expected-loss estimator introduced in this paper is, to our knowledge, novel. It should be considered an alternative method to the classic weighted kernel method (Cai, 2001; Salha & El Shekh Ahmed, n.d.). While we do not attempt a detailed theoretical study of this estimator, we do establish the need for the shape constraint in our spline estimator. Though this fact is probably well known, we prove a (slightly stronger) version here for completeness.

**Proposition 1.** *Given a collection of metric spaces* *X*_1_, …*X*_*n*_ with metrics *d*_1_…*d*_*n*_ (e.g. *d*_*centroid*_, *d*_*c*_*re), and a function f* : (*X*_1_ × *X*_1_)… × (*X*_*n*_ × *X*_*n*_) = *g* (*d*_1_(*X*_1_ × *X*_1_), …*d*_*n*_(*X*_*n*_ × *X*_*n*_)), *then* *f* *is a metric if *g* is concave, non-decreasing and* *g* (*d*) = 0 ⇔ *d* = 0.

*Proof.* We show *g* satisfying the above properties implies that *f* is a metric.

- The first property of a metric is that *f* (*x*, *x*') *=* 0 *⇔ x = x*'. The left implication:
- 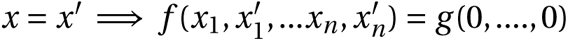, since *𝒟* are metrics. Then, since *g* (0) *=* 0, we have that *f* (*x*, *x*') *=* 0. The right implication: *f* (*x*, *x*') *=* 0 = *d* =0 *= x = x*' since *d* are metrics.
- The second property of a metric is that *f* (*x*, *x*') *= f* (*x*', *x*). This follows immediately from the symmetry of the *d_i_*, i.e. 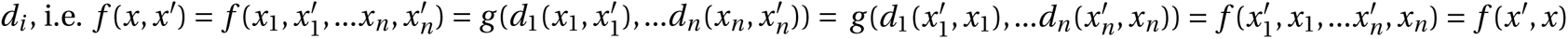
- The third property of a metric is the triangle inequality: *f* (*x*, *x*') *≤ f* (*x*, *x\**) *+ f* (*x\**, *x*'). To show this is satisfied for such a *g*, we first note that *f* (*x*, *x*') *= g* (*d* (*x*, *x*')) *≤ g* (*d* (*x*, *x\**) *+d* (*x\**, *x*')) since *g* is non-decreasing and by the triangle inequality of *𝒟* . Then, since *g* is concave, *g* (*d* (*x*, *x\**) *+d* (*x\**, *x*')) *≤ g* (*d* (*x*, *x\**)) *+g* (*d* (*x\**, *x*')) *= f* (*x*, *x\**) *+ f* (*x\**, *x*').

### Setting a lower detection threshold

The lower detection threshold of our approach is a complicated consequence of our experimental and analytical protocols. For example, the Nadaraya-Watson estimator is likely to generate many small false positive connections, since the projection of even a single experiment within the source region to a target will cause a non-zero connectivity in the Nadaraya-Watson weighted average. On the other hand, the complexities of the experimental protocol itself and the image analysis and alignment can also cause spurious signals. Therefore, it is of interest to establish a lower-detection threshold below which we have very little power-to-predict, and set estimated connectivities below this threshold to zero.

We set this threshold with respect to the sum of Type 1 and Type 2 errors

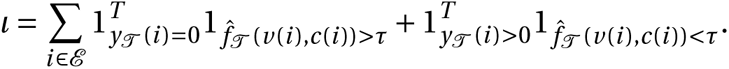

We then select the *τ* that minimizes *l*. Results for this approach are given in Supplemental Section 7.

### Decomposing the connectivity matrix

We utilize non-negative matrix factorization (NMF) to analyze the principal signals in our connectivity matrix. Here, we review this approach as applied to decomposition of the distal elements of the estimated connectivity matrix 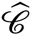 to identify *q* connectivity archetypes. Aside from the NMF program itself, the key elements are selection of the number of archetypes *q* and stabilization of the tendency of NMF to give random results over different initializations.

#### Non-negative matrix factorization

As discussed in Knox et al. (2019), one of the most basic processes underlying the observed connectivity is the tendency of each source region to predominantly project to proximal regions. For example, the heatmap in Supplemental Figure 17 shows that the pattern of intrastructure distances resembles the connectivity matrix in 2. These connections are biologically meaningful, but also unsurprising, and their relative strength biases learned latent coordinate representations away from long-range structures. For this reason, we establish a 1500*µm* ’distal’ threshold within which to exclude connections for our analysis.

Given a matrix 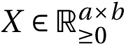 and a desired latent space dimension *q*, the non-negative matrix factorization is thus

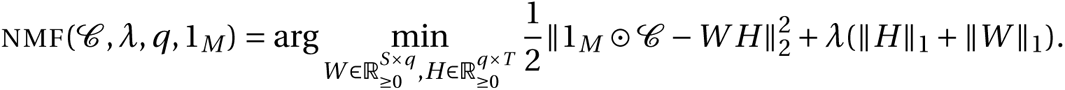

The mask 1*_M_* specifies this objective for detecting patterns in long-range connections. We note the existence of NMF with alternative norms for certain marginal distributions, but leave utilization of this approach for future work (Brunet et al., 2004).

The mask 1*_M_* ∈ {0, 1}*^S×T^* serves two purposes. First, it enables computation of the NMF objective while excluding self and nearby connections. These connections are both strong and linearly independent, and so would unduly influence the *NMF* reconstruction error over more biologically interesting or cell-type dependent long-range connections. Second, it enables cross-validation based selection of the number of retained components.

#### Cross-validating NMF

We review cross-validation for NMF following (Perry, 2009). In summary, a NMF model is first fit on a reduced data set, and an evaluation set is held out. After random masking of the evaluation set, the loss of the learned model is then evaluated on the basis of successful reconstruction of the held-out values. This procedure is performed repeatedly, with replicates of random masks at each tested dimensionality *q*. This determines the point past which additional hidden units provide no additional value for reconstructing the original signal.

The differentiating feature of cross-validation for NMF compared with supervised learning is the randomness of the masking matrix 1*_M_* . Cross-validation for supervised learning generally leaves out entire observations, but this is insufficient for our situation. This is because, given *W*, our *H* is the solution of a regularized non-negative least squares optimization problem

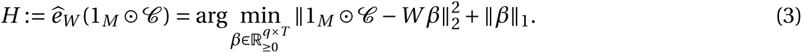

The negative effects of an overfit model can therefore be optimized away from on the evaluation set.

We therefore generate uniformly random masks 1*_M_*_(_*_p_*_)_ ∈ **ℝ***^S×T^* where

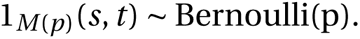

NMF is then performed using the mask 1*_M_*_(_*_p_*_)_ to get *W* . The cross-validation error is then

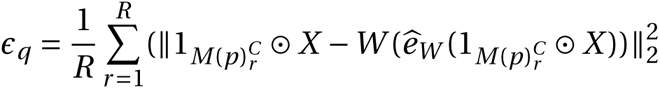

where 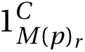 is the binary complement of 1*_M_*_(_*_p_*_)_*_r_* and *ℛ* is a number of replicates. Theoretically, the optimum number of components is then

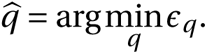

#### Stabilizing NMF

The NMF program is non-convex, and, empirically, individual replicates will not converge to the same optima. One solution therefore is to run multiple replicates of the NMF algorithm and cluster the resulting vectors. This approach raises the questions of how many clusters to use, and how to deal with stochasticity in the clustering algorithm itself. We address this issue through the notion of clustering stability (von Luxburg, 2010a).

The clustering stability approach is to generate *L* replicas of k-cluster partitions {*C_kl_* : *l* ∈ 1… *L*} and then compute the average dissimilarity between clusterings

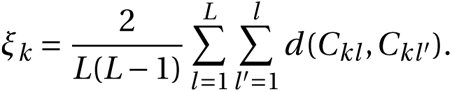

Then, the optimum number of clusters is

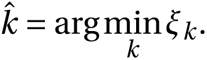

A review of this approach is found in von Luxburg (2010b). Intuitively, archetype vectors that cluster together frequently over clustering replicates indicate the presence of a stable clustering. For *𝒟*, we utilize the adjusted Rand Index - a simple dissimilarity measure between clusterings. Note that we expect to select slightly more than the *q* components suggested by cross-validation, since archetype vectors which appear in one NMF replicate generally should appear in others. We then select the *q* clusters with the most archetype vectors - the most stable NMF results - and take the median of each cluster to create a sparse representative archetype Kotliar et al. (2019); Wu et al. (2016). We then find the according *H* using Program 3. Experimental results for these cross-validation and stability selection approaches are given in Supplemental Section 7.

## 7 SUPPLEMENTAL EXPERIMENTS

The supplemental experiments show results on lower limit of detection, performance of our estimator for different regions and cell-classes, heirarchical clustering of connectivities, and stability and component analysis of our NMF results.

### Setting detection threshold τ

We give results on the false detection rate at different limits of detection. These conclusively show that 10^−6^ is the good threshold for our normalized data.

**Figure 19:**
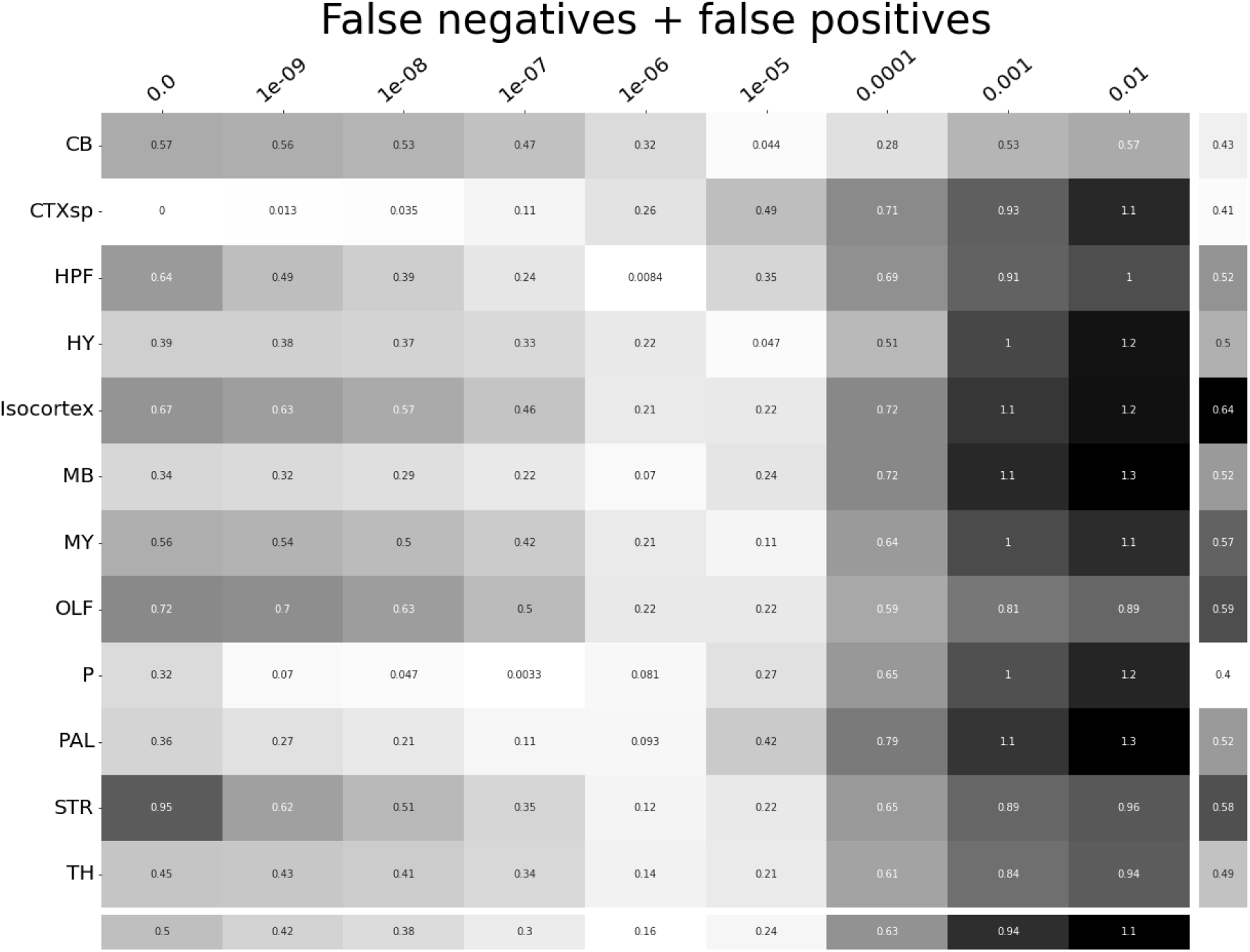
*τ* at different limits of detection in different major structures. 10^−6^ is the optimal detection threshold.

### Loss subsets

We report model accuracies for our *EL* model by neuron class and structure. These expand upon the results in Table 5 and give more specific information about the quality of our estimates. CTXsp is omitted due to the small evaluation set.

**Figure 20:**
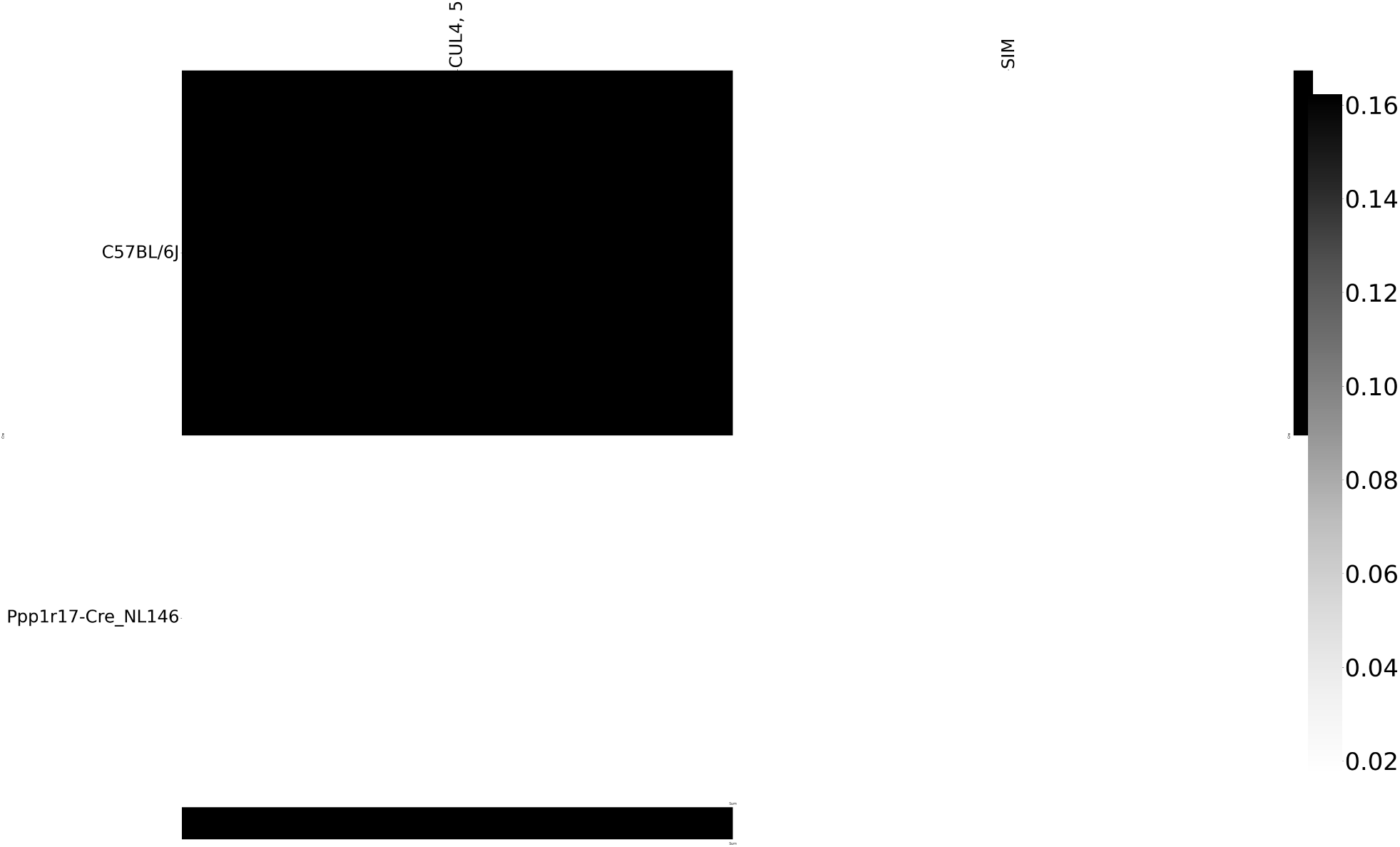
Weighted loss for Cre-leaf combinations in CB. Missing values are omitted. For example, this figure has one present and three missing values. Row and column averages are also plotted.

**Figure 21:**
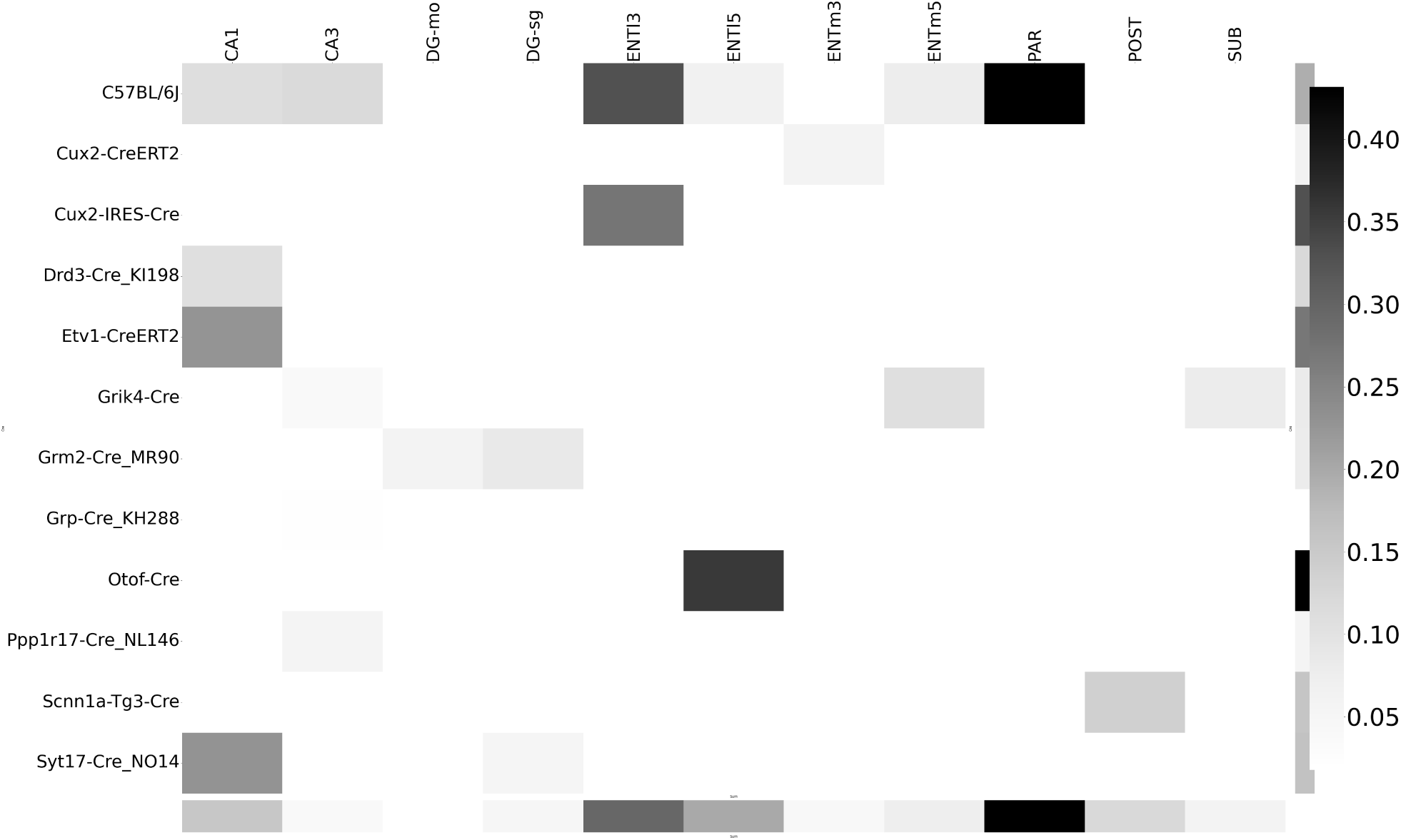
Weighted loss for Cre-leaf combinations in HPF. Missing values are omitted. Row and column averages are also plotted.

**Figure 22:**
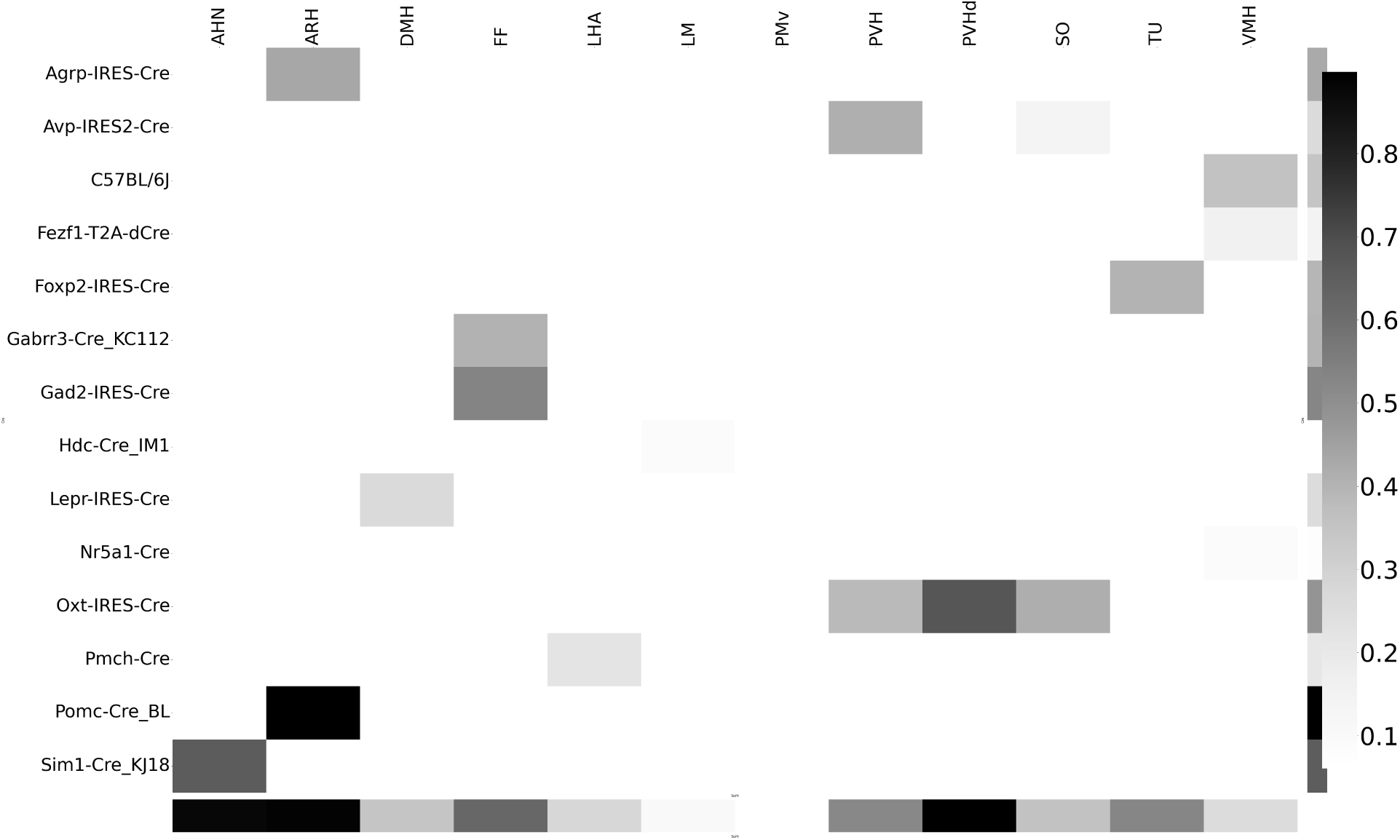
Weighted loss for Cre-leaf combinations in HY. Missing values are omitted. Row and column averages are also plotted.

**Figure 23:**
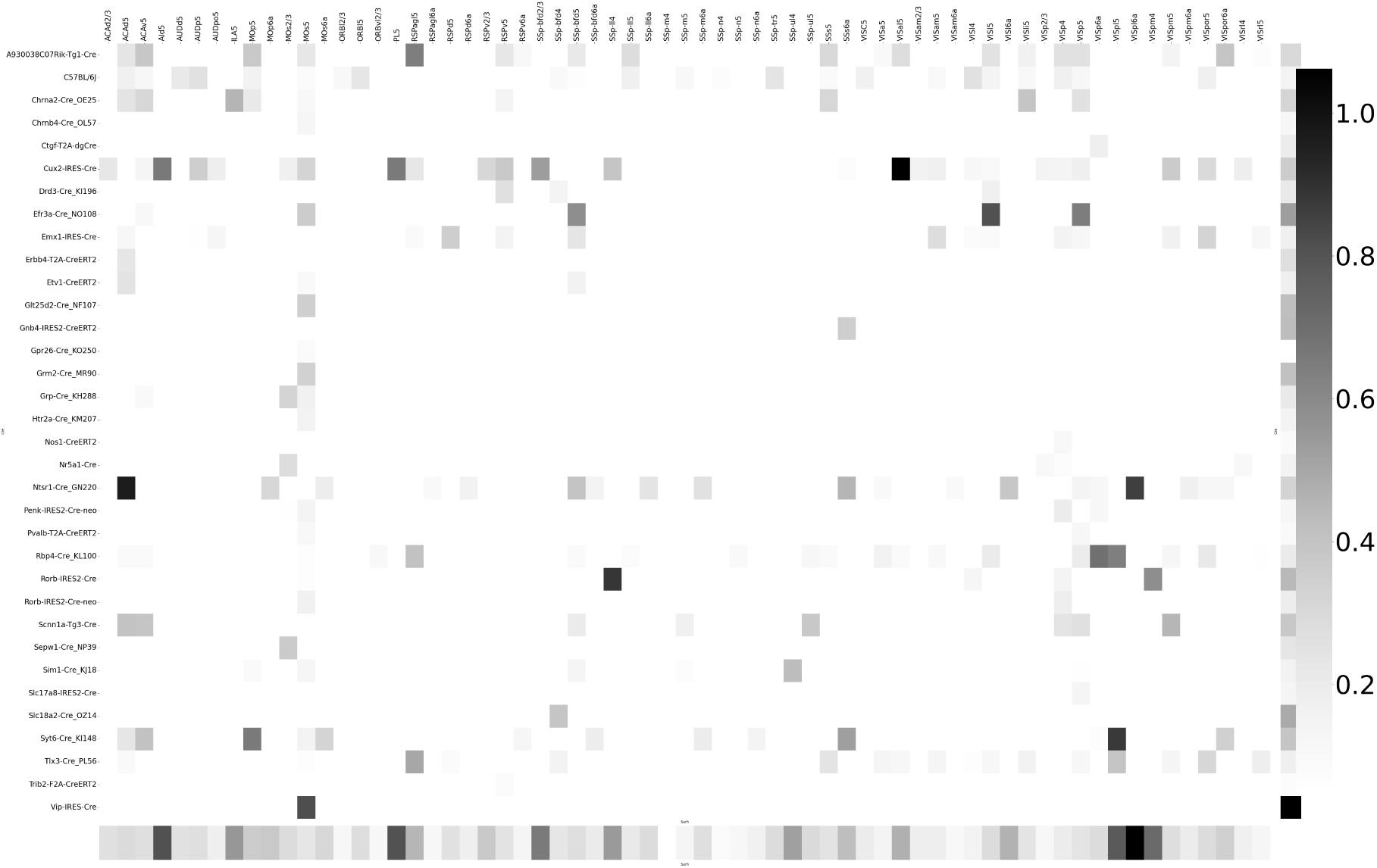
Weighted loss for Cre-leaf combinations in Isocortex. Missing values are omitted. Row and column averages are also plotted.

**Figure 24:**
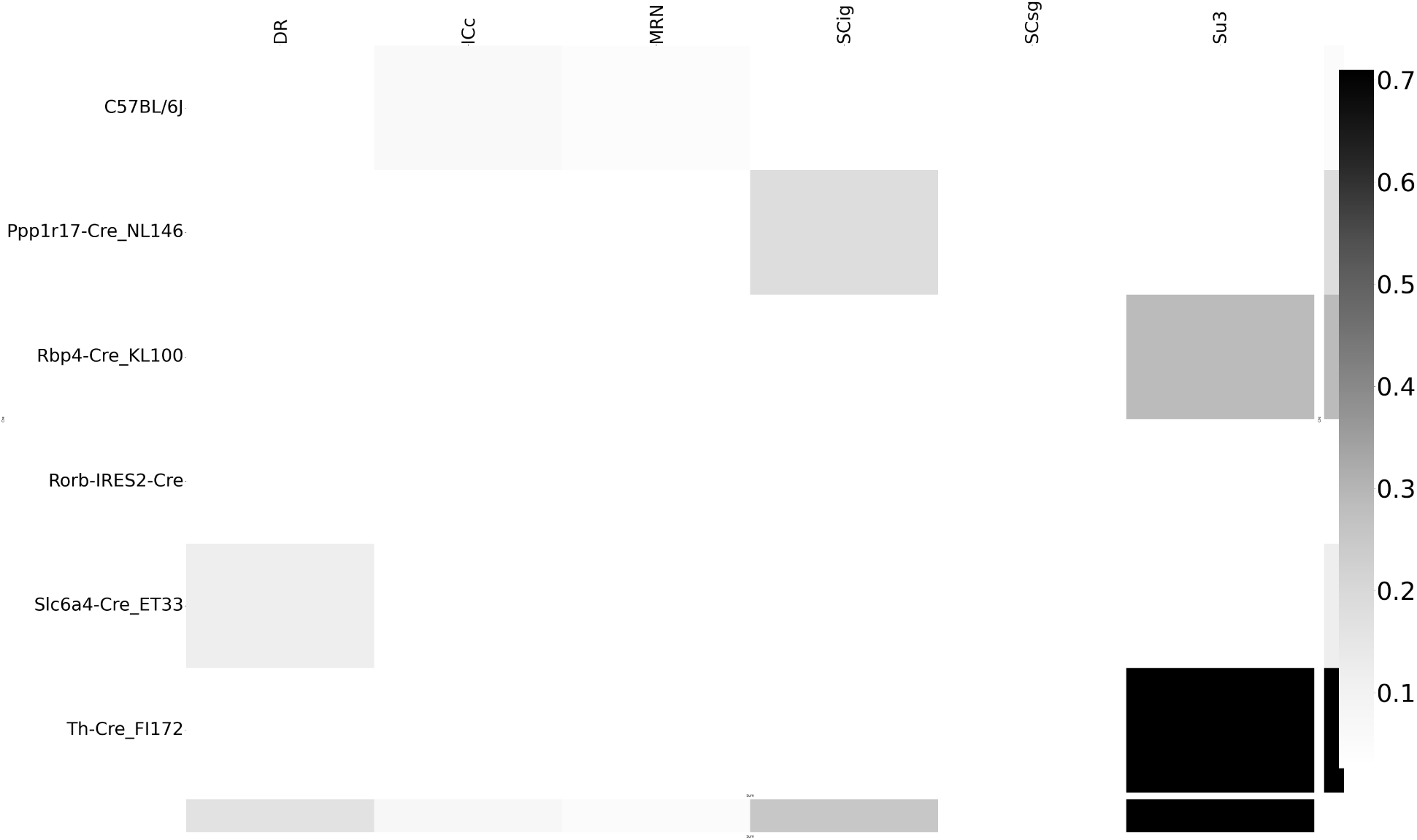
Weighted loss for Cre-leaf combinations in MB. Missing values are omitted. Row and column averages are also plotted.

**Figure 25:**
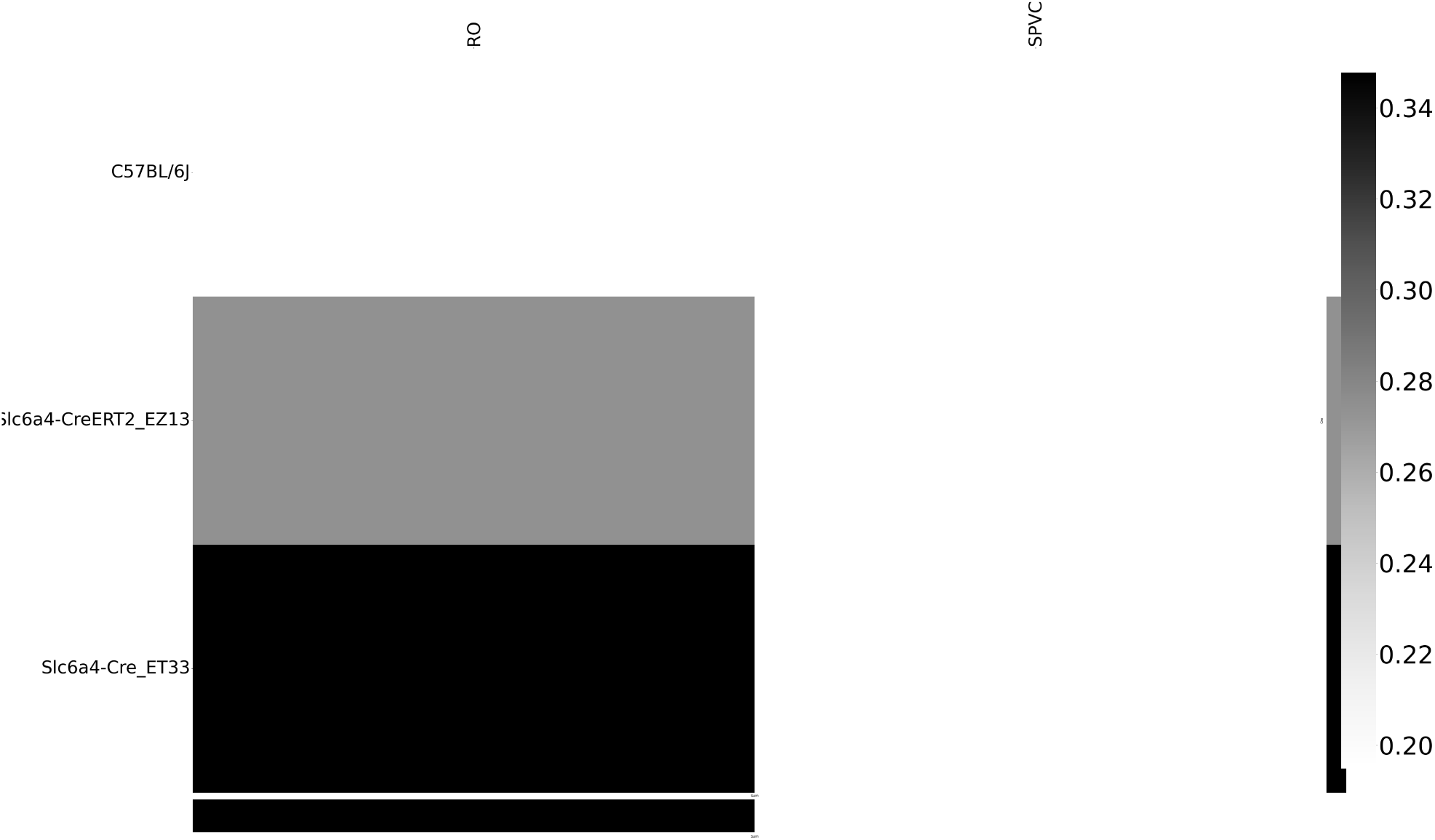
Weighted loss for Cre-leaf combinations in MY. Missing values are omitted. Row and column averages are also plotted.

**Figure 26:**
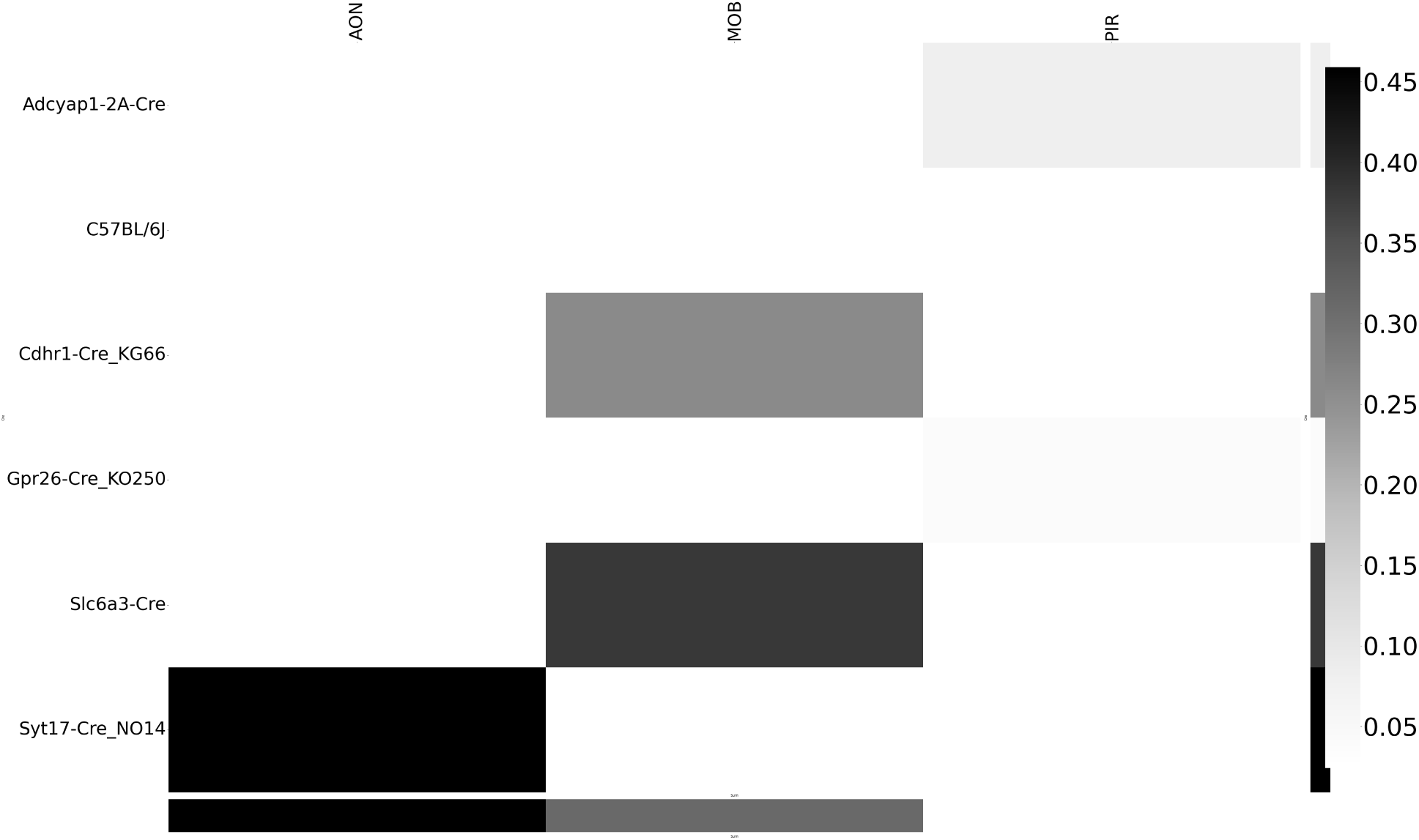
Weighted loss for Cre-leaf combinations in OLF. Missing values are omitted. Row and column averages are also plotted.

**Figure 27:**
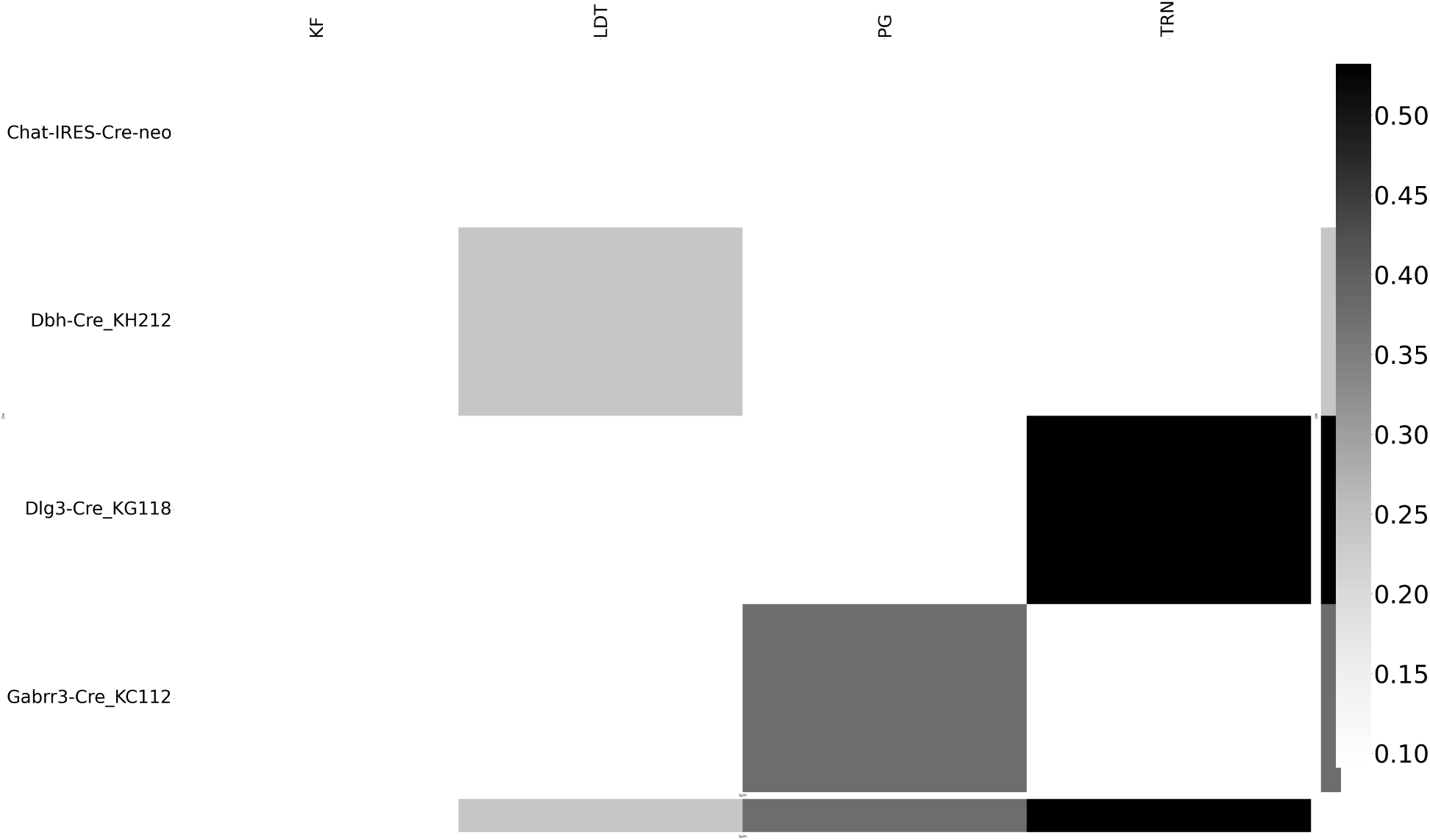
Weighted loss for Cre-leaf combinations in P. Missing values are omitted. Row and column averages are also plotted.

**Figure 28:**
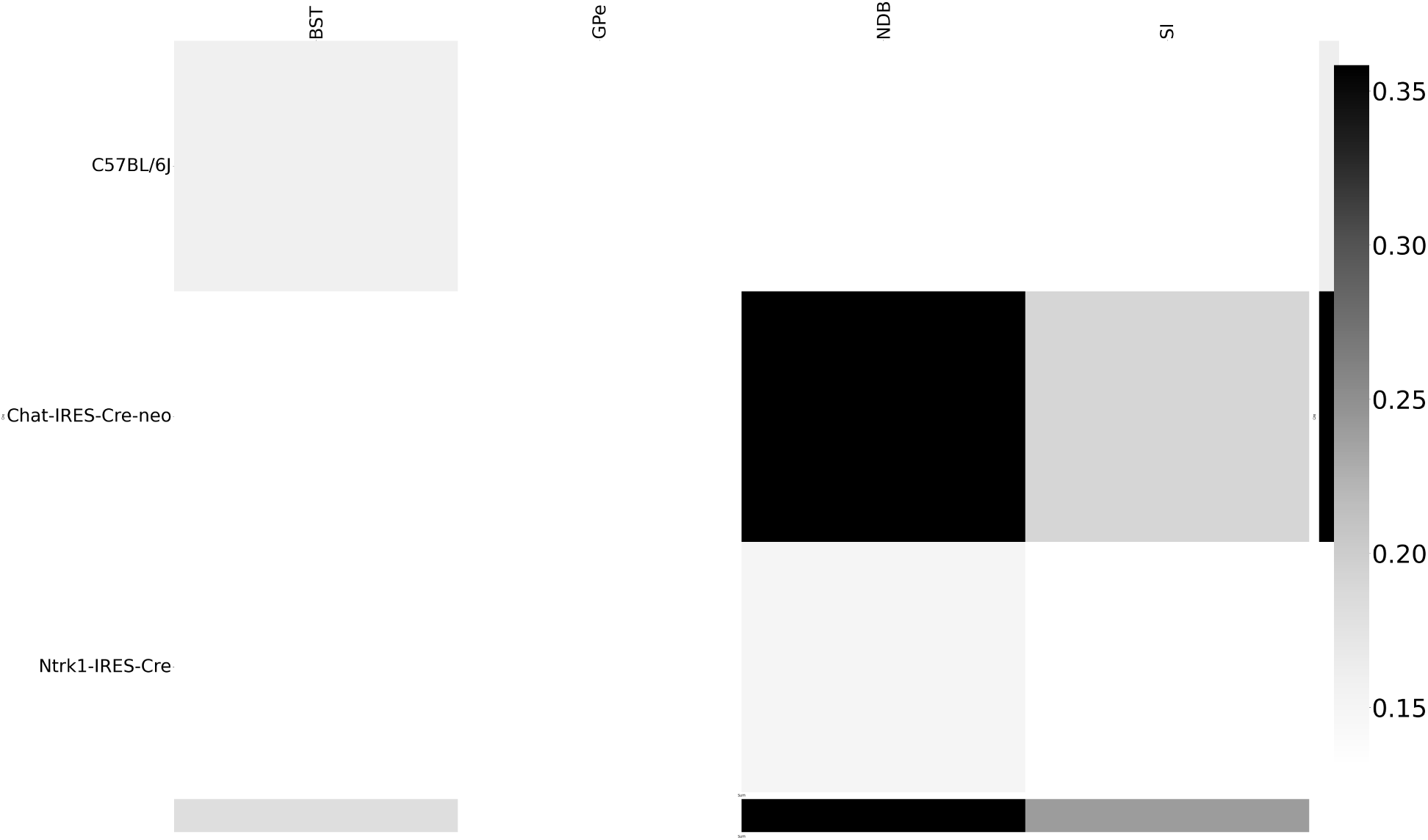
Weighted loss for Cre-leaf combinations in PAL. Missing values are omitted. Row and column averages are also plotted.

**Figure 29:**
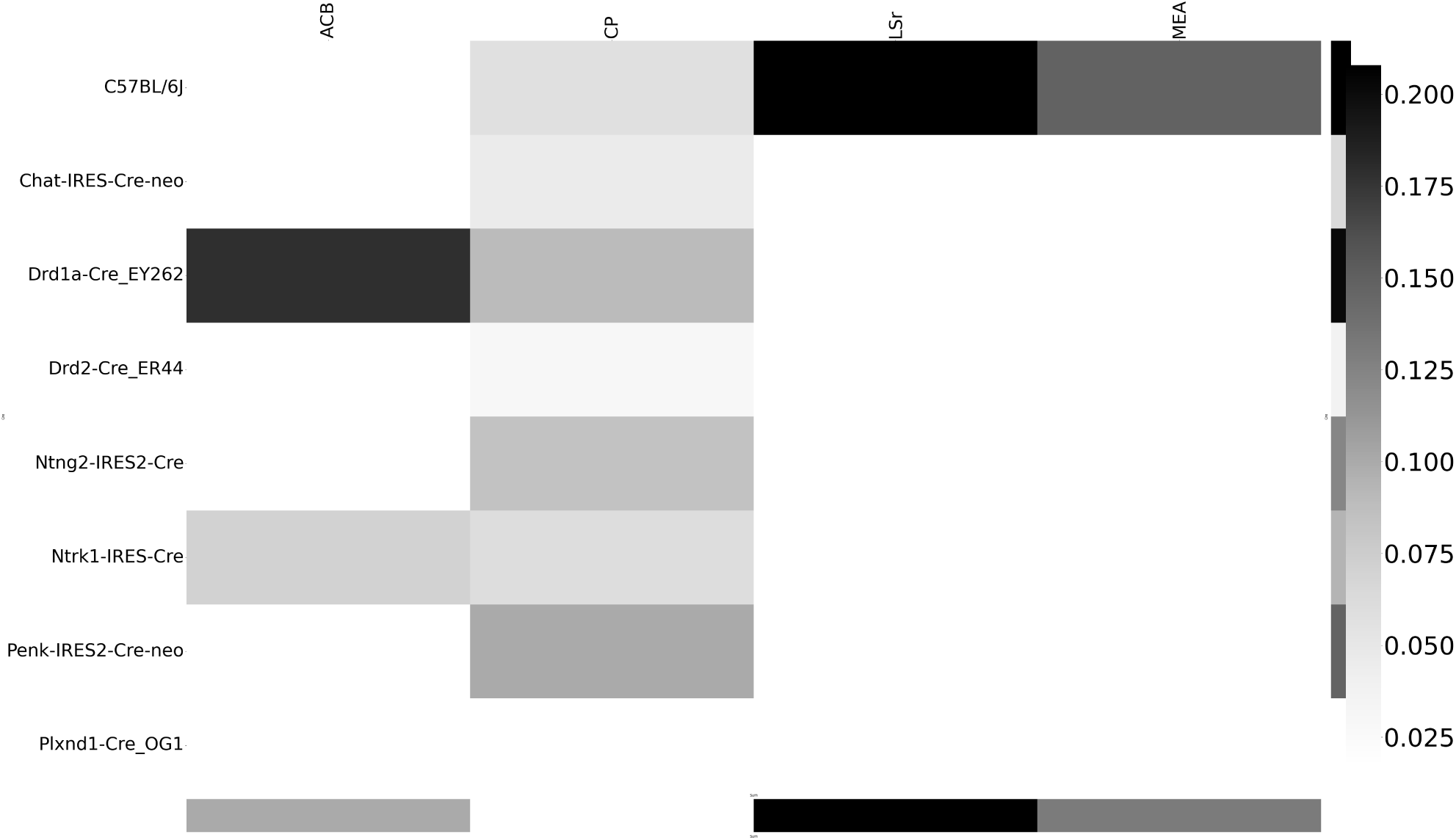
Weighted loss for Cre-leaf combinations in STR. Missing values are omitted. Row and column averages are also plotted.

**Figure 30:**
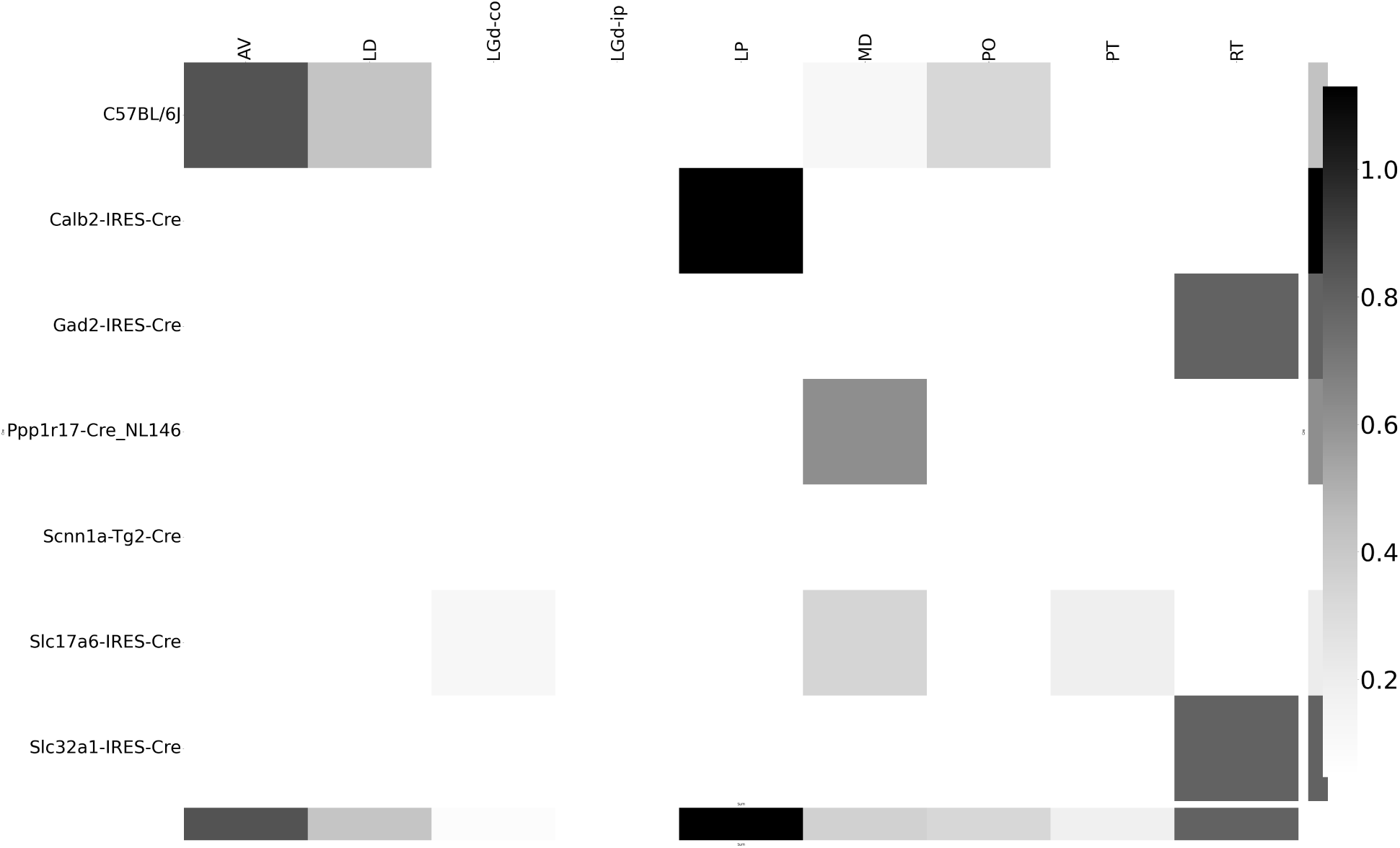
Weighted loss for Cre-leaf combinations in TH. Missing values are omitted. Row and column averages are also plotted.

### Cell-type specificity

We performed heirarchical clustering using the default method in Seaborn (Waskom, 2021) to investigate shared projection patterns across Cre-lines. That is, we used agglomerative clustering with Ward’s criterion Hastie et al. (n.d.); Lalloué et al. (2013). This showed clustering of Ntsr1 projections to Thalamic nuceli.

**Figure 31:**
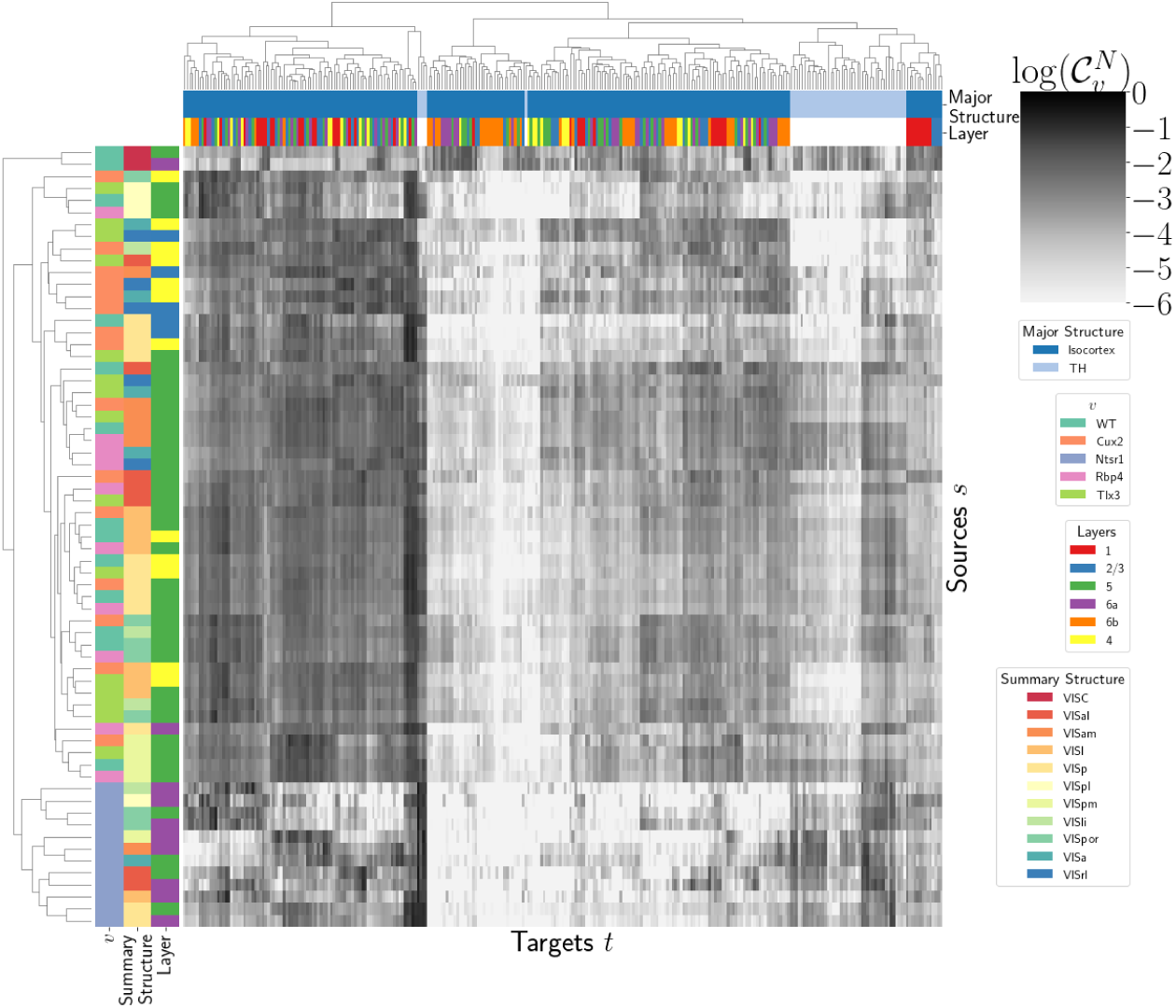
Hierarchical clustering of connectivity strengths from visual cortex cell-types to cortical and thalamic targets. Cre-line, summary structure, and layer are labelled on the sources. Major brain division and layer are labelled on the targets.

### Matrix Factorization

We give additional results on the generation of the archetypal connectome latent variables. These consist of cross-validation selection of *q*, the number of latent components, stability analysis, and visualization of the reconstructed wild-type connectivity.

#### Cross-validation

We set α *=* 0.002 and run Program 2 on *𝒷_wt_* . We use a random mask with *p =* .3 to evaluate prediction accuracy of models trained on the unmasked data on the masked data. To account for stochasticity in the NMF algorithm, we run *R =* 8 replicates at each potential dimension *q*. The lowest mean test error was observed at 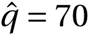 *=* 70, indicating that even more components could be estimated. However, the low decrease in reconstruction error at higher values of *q* and need for brevity in our figures motivated us to choose *q =* 15 for the purposes of display.

**Figure 32:**
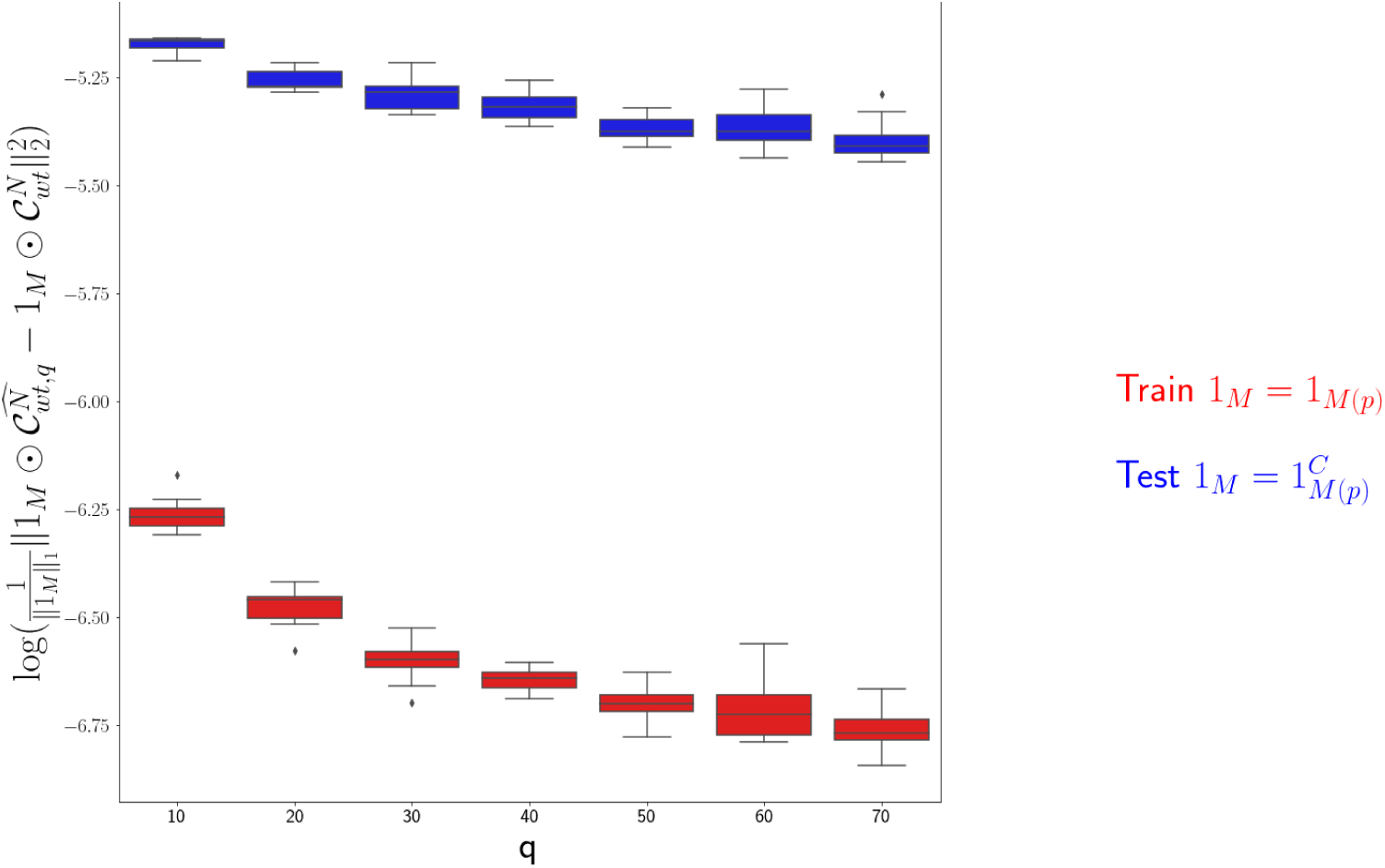
Train and test error using NMF decomposition.

#### Stability

To address the instability of the NMF algorithm in identifying components, we k-means cluster components over *R =* 10 replicates with *k* ∈ {10, 15, 20, 25, 30}. Since the clustering is itself unstable, we repeat the clustering 25 times and select the *k* with the largest Rand index, a standard method of clustering stability (Meila, 2007; Rand, 1971).

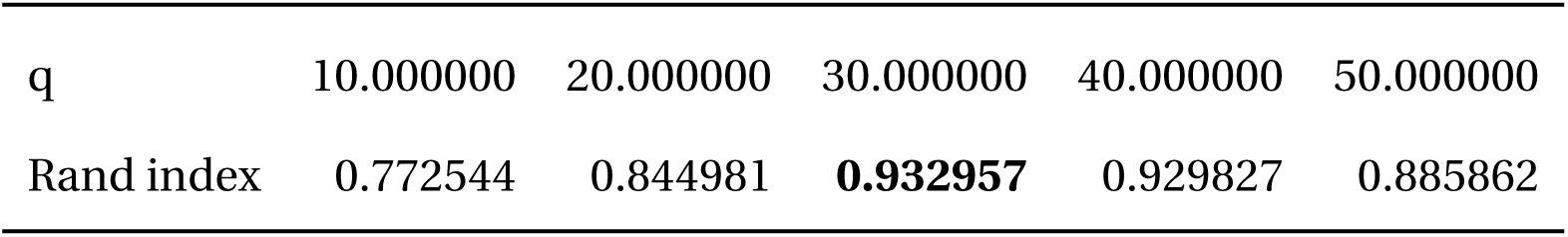

Since *k*-means is most stable at *k =* 30, we cluster the *qR =* 150 components into 30 clusters and select the 15 clusters appearing in the most replicates.

**Figure 33:**
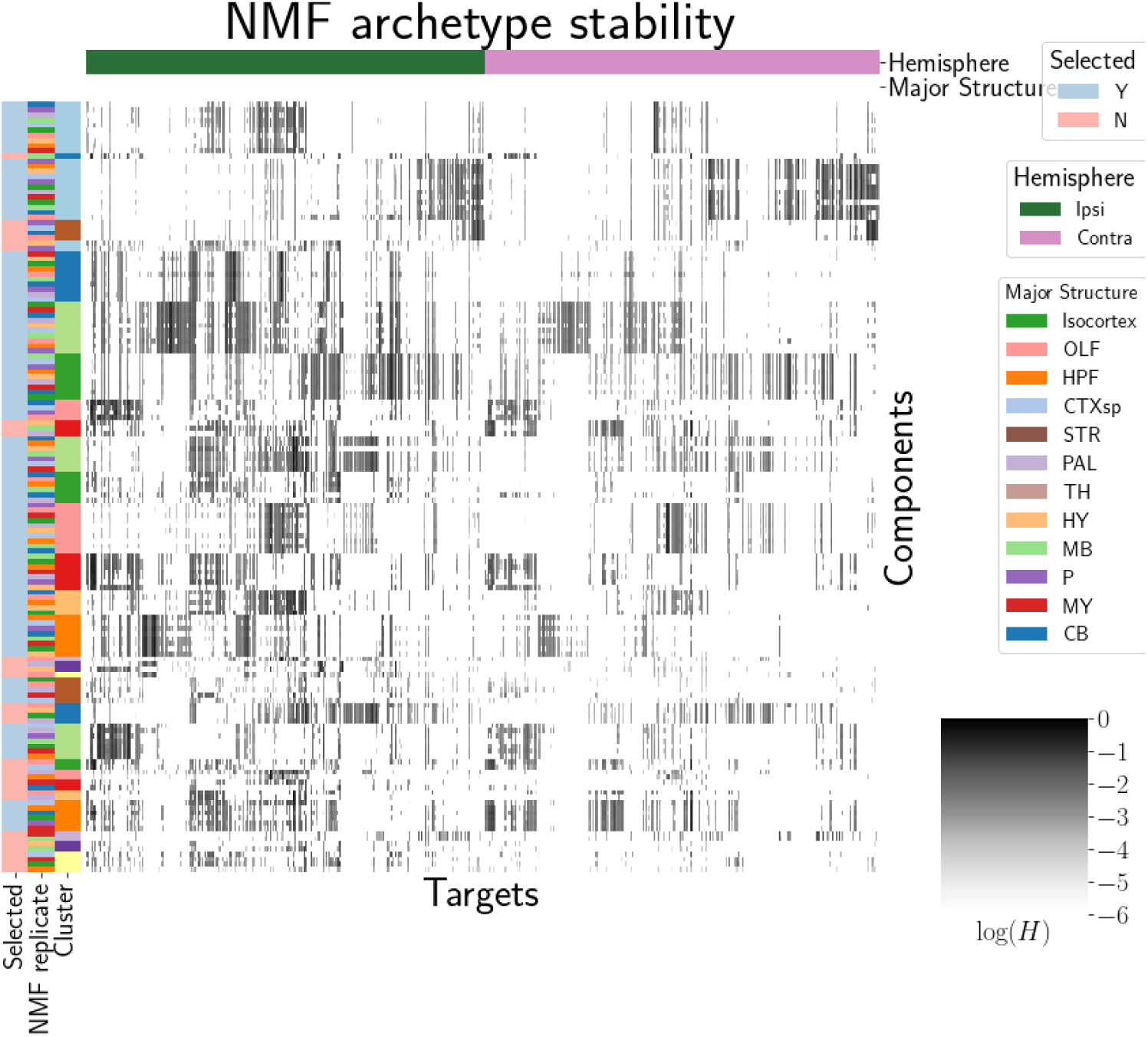
Stability of NMF results across replicates. Replicate and NMF component are shown on rows. Components that are in the top 15 are also indicated.

We plot the medians of these components in Figure 4a and in the main text. These are the connectivity archetypes. We then fit a non-negative least squares (the second step in the standard NMF optimization algorithm) to determine *W* (Lee & Seung, 2000)

#### Association with Cre-line

Finally, we show the association of our learned archetypes with projections from sources with injection centroids from the Ntsr1, Cux2, Rbp4, and Tlx3 Cre-lines. While we make no statistical claims on these associations, the distribution of cosine similarities of sources from each of the Cre-line lines shows an association of learned archetypes with Cre-line.

**Figure 34:**
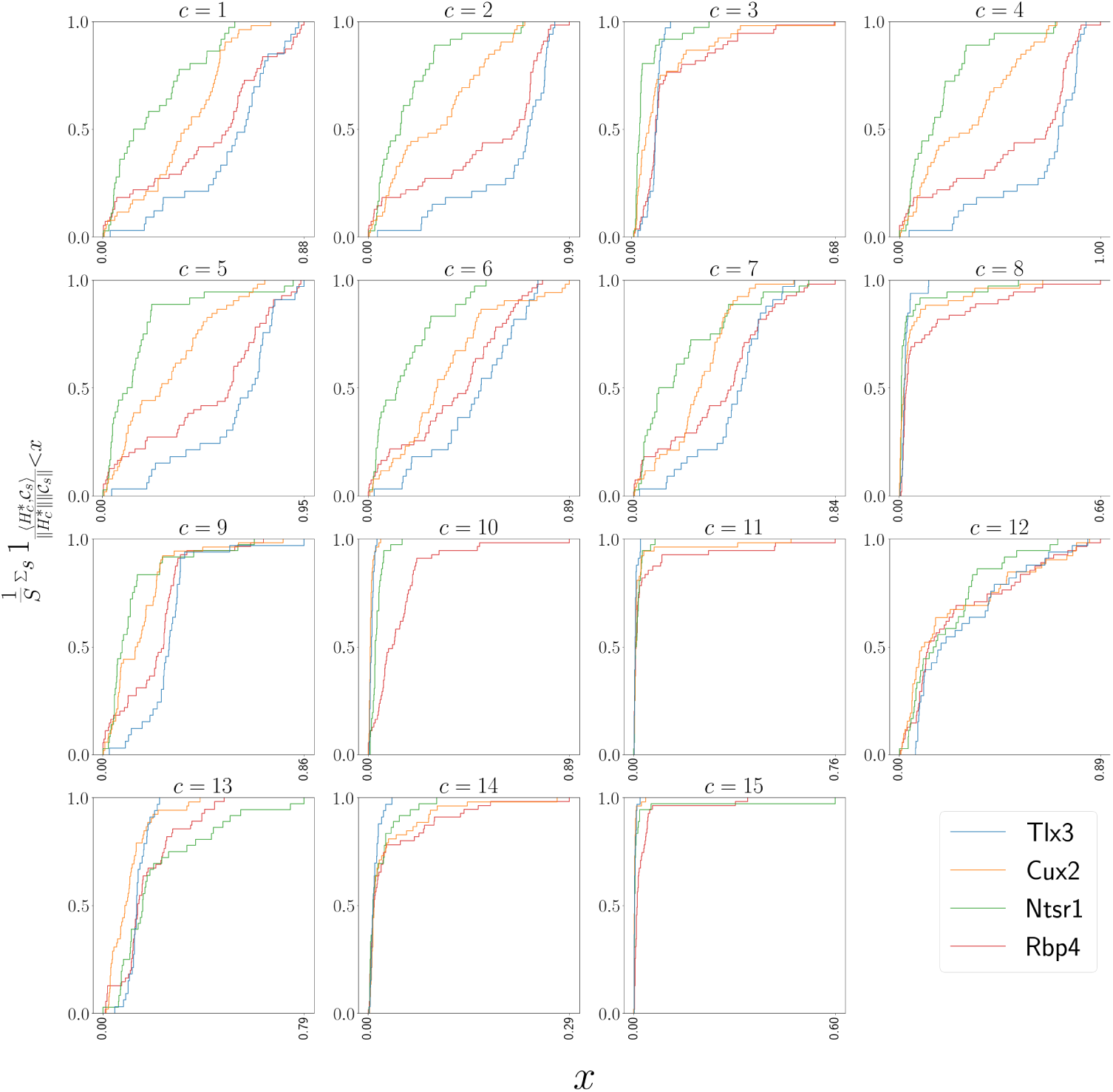
Empirical cumulative distributions of cosine similarities between source structures and connectivity components for four different Cre-lines.

## 8 TECHNICAL TERMS

**Technical Term** a key term that is mentioned in an NETN article and whose usage and definition may not be familiar across the broad readership of the journal.

**Cre-line** The combination of Cre-recombinase expression in transgenic mouse and Cre-induced promotion in the vector that induces labelling of projection

**Cell class** The projecting neurons targeted by a particular Cre-line **Structural connectivities** connectivity between structures

**Voxel** A 100*µm* cube of brain

**Structural connection tensor** Connectivities between structures given a neuron class

**dictionary-learning** A family of algorithms for finding low-dimensional data representations.

**Shape constrained estimator** A statistical estimator that fits a function of a particular shape (e.g. monotonic increasing, convex).

**Nadaraya-Watson** A simple local smoothing estimator.

**Connectivity archetypes** Projection patterns from which we can linearly reconstruct the connectome

**Expected loss** Our new estimator that weights different features by their estimated predictive power.

